# Uncovering brain tissue architecture across scales with super-resolution light microscopy

**DOI:** 10.1101/2022.08.17.504272

**Authors:** Julia M. Michalska, Julia Lyudchik, Philipp Velicky, Hana Korinkova, Jake F. Watson, Alban Cenameri, Christoph Sommer, Alessandro Venturino, Karl Roessler, Thomas Czech, Sandra Siegert, Gaia Novarino, Peter Jonas, Johann G. Danzl

## Abstract

Mapping the complex and dense arrangement of cells and their connectivity in brain tissue demands nanoscale spatial resolution imaging. Super-resolution optical microscopy excels at visualizing specific molecules and individual cells but fails to provide tissue context. Here we developed Comprehensive Analysis of Tissues across Scales (CATS), a technology to densely map brain tissue architecture from millimeter regional to nanoscopic synaptic scales in diverse chemically fixed brain preparations, including rodent and human. CATS leverages fixation-compatible extracellular labeling and advanced optical readout, in particular stimulated-emission depletion and expansion microscopy, to comprehensively delineate cellular structures. It enables 3D-reconstructing single synapses and mapping synaptic connectivity by identification and tailored analysis of putative synaptic cleft regions. Applying CATS to the hippocampal mossy fiber circuitry, we demonstrate its power to reveal the system’s molecularly informed ultrastructure across spatial scales and assess local connectivity by reconstructing and quantifying the synaptic input and output structure of identified neurons.

## Introduction

The challenge of illuminating the complex structure of brain tissue has been a major motivating force to advance imaging technologies. Optical super-resolution approaches visualize cells and molecules down to nanoscopic scales, increasing resolution beyond the diffraction limit of a few hundred nanometers either by increasing instrument resolution^1–4^ or by physically increasing specimen size and hence distances between features^5–8^. Super-resolution microscopy has generated insights into the molecular organization of synapses^9–11^, the neuronal cytoskeleton^12^, structure-function relationships in neurons^13^, and tissue organization^14^. However, in all these cases, analysis is limited to specific molecular targets or sparse subsets of labeled cells, lacking information about their context within the tissue. Electron microscopy (EM) provides comprehensive structural contrast and exquisite spatial resolution, but 3D-tissue reconstruction is technically challenging, laborious, and difficult to complement with molecular information. Optical technologies that visualize the tissue’s architecture and provide contextual meaning to molecules and cellular structures at high resolution would provide major opportunities for discovery.

Extracellular labeling is a powerful tool to delineate all cells in a tissue volume. It has been applied to guide patch clamp experiments^15^ and visualize extracellular space^16,17^ in living brain tissue, and for early EM-connectomics studies in mouse retina^18^. Reading out freely diffusing, extracellularly applied fluorophores with stimulated emission depletion (STED) microscopy^1,19,20^ in living brain tissue in the framework of super-resolution shadow imaging (SUSHI)^17,21–23^ casts super-resolved shadows of all cells. We recently showed that extracellular labeling integrated with a specifically engineered 3D-super-resolution/machine learning technology enables dense, synapse-level reconstruction of living brain tissue^24^. However, while live imaging uniquely accesses dynamics, it is constrained by applicable super-resolution modality, molecular labeling options, addressable tissue volumes, and sample type. In fixed tissues, feature-rich representations of cells and various tissues have been achieved by several strategies, including fluorescent^25–29^ or Raman^30^ contrast for total protein density or other molecule classes in expansion microscopy (ExM). However, none of these approaches has been amenable to *in silico* reconstruction of the brain tissue’s architecture or sub-cellular morphology. There is thus an unmet need for an optical technology that is capable of visualizing and quantifying tissue organization from regional to single-synapse level in a technologically straightforward manner.

Here we developed *Comprehensive Analysis of Tissues across Scales* (CATS), an integrated labeling, optical imaging, and analysis platform to decode brain tissue architecture, subcellular morphologies, and molecular arrangements within their structural context. We engineered CATS to visualize all cellular structures in fixed tissue by extracellular labeling with (super-resolution) fluorescence microscopy. Thereby, CATS removes the constraints associated with live imaging and permits analysis from regional to nanoscopic spatial scales for commonly used native and cultured brain tissue preparations. It capitalizes on the full technology base for labeling, optically homogenizing, and 3D-super-resolution imaging available for fixed tissues, building on the pertinent strengths of STED and expansion microscopy, in a widely adoptable approach. With specifically tailored analysis, CATS quantitatively reveals tissue architecture, maps synaptic connectivity, and allows 3D-reconstruction of subcellular morphology down to single-synapse level in a molecularly informed fashion. To demonstrate the power of this approach to quantify synaptic connectivity and structure, we characterized one of the key synapse types in the hippocampal circuitry. We reveal the synaptic input and output structure of identified, functionally recorded neurons across brain regions, and furthermore apply the technique to clinically derived human tissue samples.

## Results

### CATS unravels tissue architecture at super-resolved detail

We developed two strategies for revealing tissue structure by selective labeling of the extracellular domain (**Fig. 1a**): i) “Compartment CATS” (coCATS) applies covalently binding labeling compounds to the *extracellular compartment* in living tissue, with intact membrane boundaries constraining labeling to the extracellular space and cell surfaces. ii) “Resident CATS” (rCATS) labels classes of *extracellularly resident* molecules, in particular extracellular polysaccharides. This makes CATS applicable also to specimens where live labeling is not possible (**Fig. 1b**). Both approaches revealed the cellular architecture of brain tissue, for example in the hippocampal region, across scales (**Fig. 1b,c**).

**Fig. 1.**
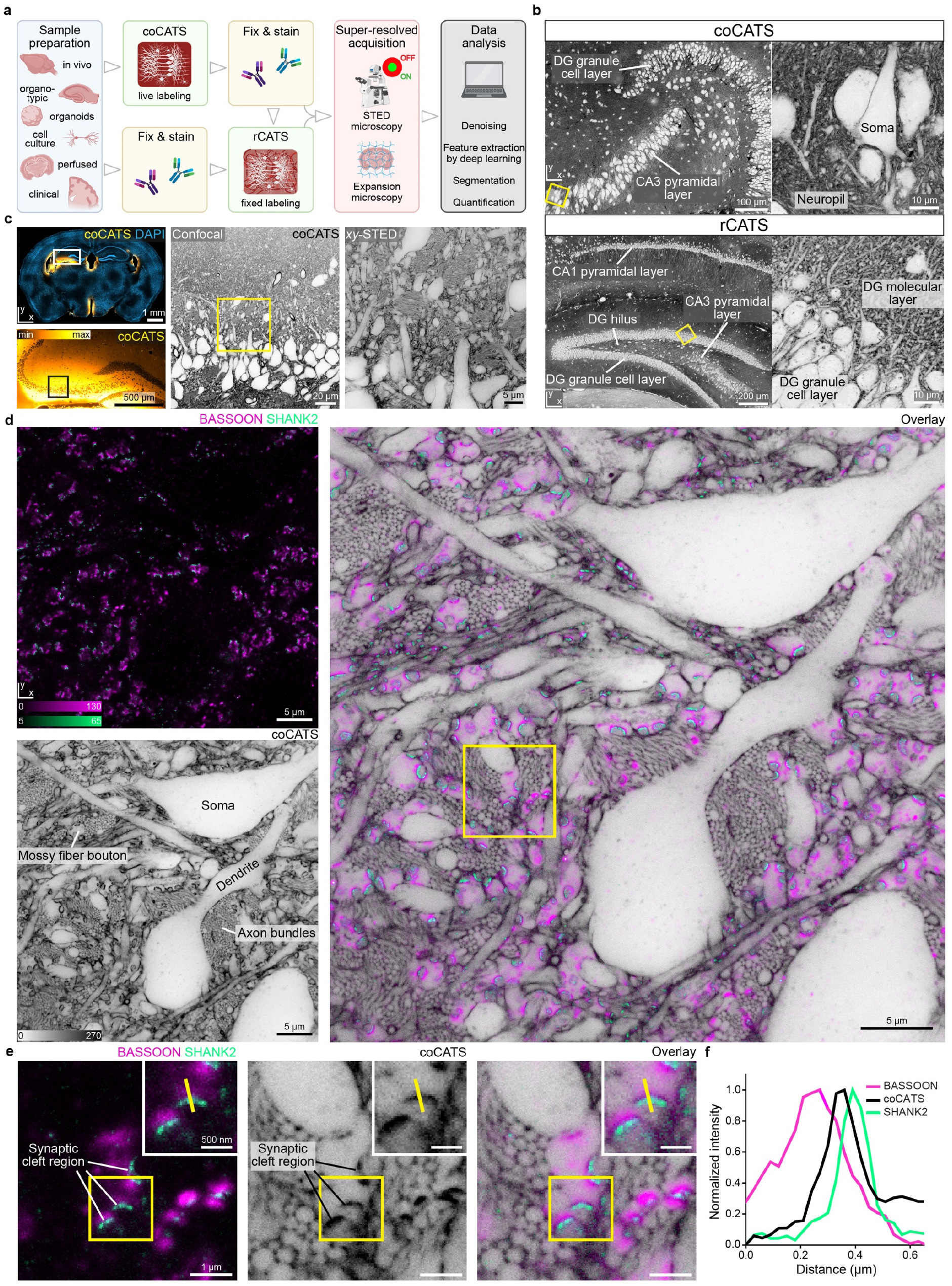
Comprehensive analysis of tissue across scales (CATS). **a**, Platform for tissue analysis including live extracellular labeling (compartment CATS, coCATS) or extracellular labeling in previously fixed tissue (resident CATS, rCATS), optional molecular staining, super-resolved acquisition, and machine learning/conventional analysis. **b**, (*Top*) CoCATS labeling in organotypic hippocampal slice, revealing gross architecture of the dentate gyrus (DG) and CA3 region, and zoomed view of boxed region (confocal). (*Bottom*) RCATS labeling in perfusion-fixed adult mouse coronal section, showing hippocampus with zoomed view. Intensity lookup tables (LUTs) for CATS are inverted throughout, i.e. black regions correspond to high labeling intensity, unless otherwise noted. Raw data. **c**, Progressive zoom from hippocampal regional to cellular scale in CA3 *stratum pyramidale* and *stratum lucidum*. CoCATS labeling by *in vivo* stereotactic injection into the lateral ventricle (LV) of adult mouse (*left*: LUT not inverted, *left bottom*: gamma correction applied). *Left, center*: confocal; *right*: STED, lateral resolution increase (*xy*-STED). Raw data. **d**, Super-resolved tissue architecture of mouse CA3 *stratum pyramidale/lucidum*, after *in vivo* coCATS label microinjection into LV. (*Left top*) Immunostaining of pre-synaptic BASSOON (magenta, confocal) and post-synaptic SHANK2 (turquoise, *xy*-STED). (*Left bottom*) coCATS (*xy*-STED) of same region. (*Right*) Overlay placing synaptic markers into structural context, including mossy fiber boutons (MFBs). Raw data. **e**, Magnified view from d (boxed), focusing on a MFB with multiple synaptic sites, amidst bundles of thin mossy fiber axons. Inset: magnification of synaptic transmission site. High-intensity coCATS labeling pinpoints dense/protein-rich features between pre- and post-synapses corresponding to putative synaptic cleft regions (pSCRs). **f**, Line profile as indicated in e, showing sandwich arrangement of BASSOON, high-intensity coCATS (pSCR), and SHANK2 signals.

To analyze brain architecture with coCATS, we screened for molecules providing high extra-to intracellular contrast, high labeling density, and compatibility with downstream super-resolution readout in organotypic hippocampal slice cultures (**Supplementary Fig. 1**). We focused on commercially available compounds for easy adoptability. We ensured cell impermeability either via hydrophilic, anionic fluorophores or additional sulfo- or polyethylene glycol (PEG) groups. As expected, chemistries targeting primary amines, including N-hydroxysuccinimide (NHS) and tetra-(TFP) and pentafluorophenyl (PFP) esters, effectively mediated homogeneous labeling by covalent attachment to extracellular and cell surface molecules, particularly proteins. For readout, we used either directly conjugated fluorophores or a small molecule reporter (biotin/fluorescent avidin).

For decrypting near-natively preserved brain at super-resolved detail, we performed *in vivo*-stereotactic injection of an NHS-derivative of a hydrophilic, far-red, high-performance STED fluorophore followed by transcardial fixative perfusion. Injection into the brain lateral ventricle labeled areas adjacent to the ventricular system, distant from the lesioned region at the injection site (**Supplementary Fig. 2**). STED microscopy provides direct, “all optical” super-resolved readout by applying a light pattern that confines fluorescence ability to volumes smaller than the diffraction limit. We first focused on hippocampus, a brain region central to spatial navigation and memory with well-characterized fundamental circuitry. As an important component, mossy fibers originating from dentate gyrus (DG) granule cells convey excitatory input to pyramidal neurons in the CA3 *stratum lucidum*, forming key synapses in the hippocampal trisynaptic circuit. These synapses are an established model for functional synapse characterization and contribute to higher order computations^31,32^. STED imaging at the transition between *stratum pyramidale* and *stratum lucidum* of the CA3 region revealed the complex arrangement of cell bodies, dendrites, bundles of thin axons, and synaptic terminals at high signal-to-noise ratio (SNR) (**Fig. 1d**). Diffraction-unlimited resolution, here on the order of 60 nm laterally, was indispensable to resolve the cellular structures in this extremely complex and dense arrangement (**Supplementary Fig. 3**). For example, we were able to resolve the individual unmyelinated axons in mossy fiber bundles, most conspicuous as small circular structures when transversely optically sectioned. When complemented with immunolabelling for the pre-synaptic marker BASSOON and for SHANK2, a scaffolding protein in post-synaptic densities of excitatory synapses, CATS assigned molecularly defined synaptic sites to individual pre-synaptic boutons of mossy fibers and revealed their location within the tissue’s ultrastructure (**Fig. 1d-f**). It revealed both pre- and post-synaptic structures, showing the complex arrangement of large mossy fiber boutons (MFBs) containing multiple transmission sites. These contact complex pyramidal neuron spines^33^ termed “thorny excrescences”. Such contextual structural meaning was missing with immunostainings alone or sparse labeling of a subset of cells by gold-standard cytosolic fluorescent protein expression (**Supplementary Fig. 4**).

### Quantifying single synapse structure and connectivity

When inspecting the combined structural/molecular data more closely, we discovered that coCATS labeling consistently produced high-intensity features sandwiched between pre- and post-synapses. These correspond to putatively primary amine/protein-rich extracellular regions at apparent synaptic transmission sites, likely reflecting the high protein density of the synaptic cleft^34^ (**Fig. 1e,f**). We clarified their spatial relationship with a range of additional synaptic molecules (SYNAPTOPHYSIN1, HOMER1, vesicular GABA transporter (VGAT), vesicular glutamate transporter (VGLUT1), N-CADHERIN) both in excitatory and inhibitory synapses and with sparsely labeled MFBs (**Supplementary Fig. 5**). Taken together, we found their location consistent with synaptic clefts. This prompted us to designate them “putative synaptic cleft regions” (pSCRs) and develop an automated pipeline for segmenting and mapping them (**Fig. 2a**). After enhancing volumetric datasets with a deep-learning-based denoising algorithm (Noise2Void^35^, N2V, **Supplementary Fig. 6,7**), we used super-resolved SHANK2 immunostaining as guide to excitatory synapses and performed locally confined thresholding to isolate high-intensity coCATS features. We classified them as pSCRs in case of apposition with pre-synaptic BASSOON (confocal) and post-synaptic SHANK2 (STED) in a triple-sandwich arrangement. This also eliminated false positive synapse identifications from unavoidable background in immunostainings (**Supplementary Fig. 8**). Finally, we performed instance segmentation of individual pSCRs, applied manual proofreading, and contextualized them by association with manually created volume segmentations of MFBs. Automated analysis substantially reduced human processing time compared to manual pSCR segmentation.

**Fig. 2.**
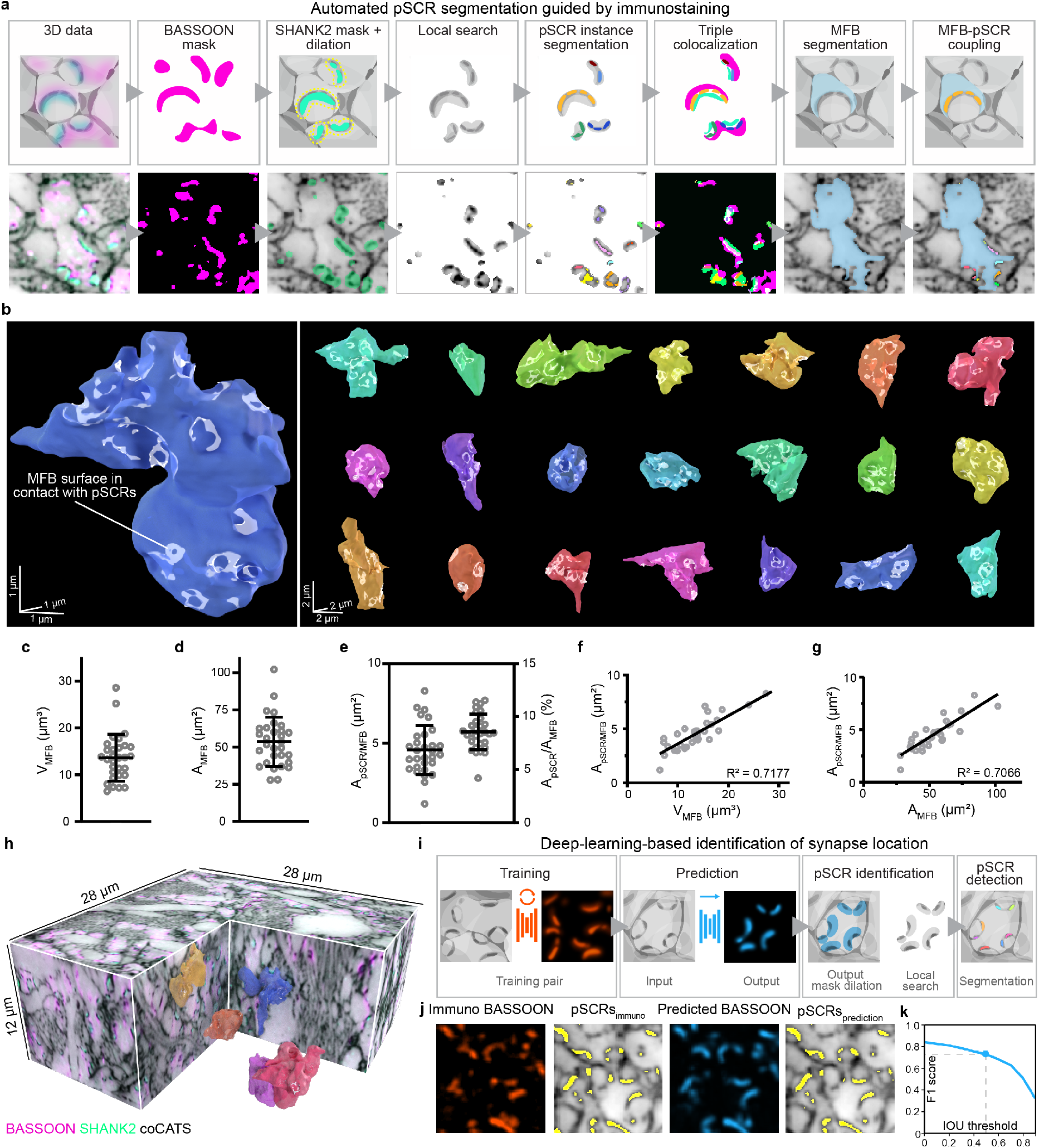
Quantifying synaptic connectivity and single-bouton properties. CoCATS analysis of hippocampal mossy fiber to CA3 pyramidal neuron synapses in adult mouse CA3 *stratum lucidum* by *in vivo* microinjection into the LV. **a**, Automated synapse detection guided by immunostaining for synaptic markers. High-intensity 3D-features in coCATS in proximity to synaptic markers are segmented and classified as pSCRs in case of triple colocalization between a pre-synaptic marker, the dense coCATS feature, and a post-synaptic marker. Detected pSCRs are then associated with manual volume segmentations of individual MFBs. Schematic (*top*) and single *xy*-planes of volumetric data (*bottom*) including coCATS (grey, *z*-STED), BASSOON (magenta, confocal) and SHANK2 (turquoise, *z*-STED) after denoising with Noise2Void (N2V, raw data: **Supplementary Fig. 7**). **b**, 3D-renderings of 22 randomly selected MFBs segmented from volumetric coCATS data at near-isotropic resolution (*z*-STED). MFB surface areas occupied by pSCRs are indicated in white. 3D-scale bars refer to bouton center. **c-e**, Quantification of MFB volume (V_MFB_), MFB surface area (A_MFB_), absolute (A_pSCR/MFB_) and relative area occupied by pSCRs on individual MFBs (A_pSCR/MFB_/A_MFB_) (n_MFB_=30). **f,g**, A_pSCR/MFB_ as a function of bouton volume and surface area with linear regression (n_MFB_=30). **h**, One of the imaging volumes used for MFB characterization (denoised with N2V, raw data: **Supplementary Fig. 7**) with coCATS (grey, *z*-STED), BASSOON (magenta, confocal) and SHANK2 (turquoise, *z*-STED), including manually segmented MFBs and automatically detected pSCRs. **i**, Deep-learning-based pSCR identification with training on paired structural (coCATS) and molecular (BASSOON immunostaining) super-resolved data. Prediction of synaptic marker location in unseen datasets is based on structural data alone. PSCRs are segmented similarly as in a, but using predicted BASSOON instead of immunostainings as guide to synaptic sites. **j**, Immunostained BASSOON (orange, *z*-STED) and BASSOON distribution predicted (blue) from coCATS structure in a dataset not included in the training. Corresponding pSCRs (yellow) segmented from coCATS data (grey, *z*-STED, N2V), guided by immunostained (pSCRs_immuno_) or predicted BASSOON (pSCRs_prediction_). **k**, Quantification of similarity between pSCRs_immuno_ and pSCRs_prediction_ by F1 score (ranging from 0 to 1, combining precision and recall) as a function of intersection over union (IOU) threshold.

Using this pipeline, we reconstructed individual synaptic boutons with their synaptic transmission topology in 3D. Reconstruction is limited by the least resolved spatial axis, i.e. along the optical (*z*-)axis. We therefore applied a dedicated light pattern for near-isotropic STED-resolution^1^ (*z*-STED, **Supplementary Fig. 3**) and recorded 3 volumetric datasets in the CA3 *stratum lucidum* (∼30×30×12 µm^3^, 2 brain slices, 1 animal). We selected 10 prominent MFBs from each, manually segmented their 3D-shape from the coCATS channel, and quantified both key geometrical parameters and pSCRs (**Fig. 2b-h, Supplementary Video 1,2, Supplementary Fig. 9**). Individual boutons varied widely in size and shape, with mean volume V̅MFB=13.6±5.0 µm^3^ (± s.d., **Fig. 2c**) and mean surface area A̅MFB=53.5±16.6 µm^2^ (**Fig. 2d**). These values are consistent with previous EM results from adult mouse^36^ (V̅MFB=13.5 µm^3^, A̅MFB=66.5 µm^2^). Mean surface area was smaller in our case as we did not include filopodial structures extending from the main bouton, as they are at the limit of the resolution used in this measurement. PSCRs were similarly diverse, often forming fenestrated structures (**Fig. 2b**). To identify regions of MFBs occupied by putative active zones, we related pSCRs to MFB segmentations. For this, we identified pSCR voxels that touched voxels of segmented MFBs, extracted the contacting surfaces and turned them into 3D-meshes. The total area of individual boutons occupied by pSCRs (ApSCR/MFB) had a mean value of A̅pSCR/MFB=4.6±1.6 µm^2^ (**Fig. 2e**). The fraction of MFB surface occupied by pSCRs (ApSCR/MFB/AMFB) calculated at individual bouton level displayed smaller spread, hinting towards a correlation between MFB size and the extent of synaptic release sites. Indeed, when plotting ApSCR/MFB as a function of MFB volume (**Fig. 2f**) (Pearson correlation coefficient r=0.844, 95% confidence interval (CI): 0.694-0.923, p<0.0001, R^2^=0.72, 30 MFBs) or surface area (**Fig. 2g**) (r=0.841, CI: 0.689-0.922, p<0.0001, R^2^=0.71, 30 MFBs), we found strong correlation, indicating that larger MFBs also engage more extensively in synaptic contacts. This is in accordance with previous studies showing a linear relationship between MFB volume and active zone extent both in organotypic slice cultures and *in vivo*^37^. Interestingly, the fraction of MFB surface area occupied by pSCRs (8.6±1.7 %) was consistent with previous quantifications of MFB surface area occupied by active zones in serial sectioning EM in adult rat on a smaller number of MFBs (9.7%)^38^. The number of pSCRs showed large variability between individual boutons (3-28, mean 13.03±5.93), similar as in an EM study on adult rat^38^, and also correlated with bouton size (**Supplementary Fig. 9**). These data demonstrate that CATS can identify synaptic transmission sites and deliver quantitative biological data at single synapse level. Our data are consistent with EM-reconstructions^36,38,39^ but include molecular information and can be obtained without complex sample preparation or serial sectioning procedures and with high experimental throughput (imaging time for 3-channel measurement per volume: ∼1.5 h).

### Deep-learning-based prediction of synapse location

Based on the prominence of pSCRs in coCATS data, we hypothesized that coCATS may reveal synapse location based on local tissue structure. We thus trained a convolutional neural network with U-net architecture^40^ to predict synapse location purely from CATS structural data using deep-learning-based image translation (**Fig. 2i, Supplementary Fig. 10**). For training, we provided the network with near-isotropically super-resolved coCATS data paired with immunostainings as molecular ground truth. We used the resulting model to predict molecule location in unseen datasets. Indeed, a model trained on coCATS and super-resolved BASSOON, present in excitatory and inhibitory synapses, was capable of guiding segmentation of pSCRs in mossy fiber boutons. The network prediction can replace the immunostainings in the pipeline above for pSCR segmentation. This is remarkable, as thresholding alone, neglecting local context, was insufficient to identify pSCRs among dense CATS features. For validation, we correlated predicted BASSOON signal with immunolabeled BASSOON in datasets not included in the training (**Supplementary Fig. 10**, Pearson correlation, r=0.818). In addition to voxel-based correlation, we evaluated resulting automated pSCR segmentation guided by immunostaining vs. segmentation guided by predicted BASSOON signal using the F1 score, and found a high degree of similarity (F1=0.73 at intersection over union (IOU) threshold 0.5, **Fig. 2j,k, Supplementary Fig. 10**). For this, instance segmentations of our deep-learning-prediction and the corresponding ground-truth dataset were compared by identifying object-pairs and calculating their IOUs (ratio of overlapping volume vs. combined volume). The F1 score as a function of IOU takes into account the precision (number of correctly segmented objects divided by number of all segmented objects) and recall (number of correctly segmented objects divided by number of all true objects). As expected, network predictions improved with the higher precision of super-resolved compared to confocal molecular signals as input data for training (**Supplementary Fig. 10**). Denoising with Noise2Void had a minor effect on prediction outcome (**Supplementary Fig. 10**). These data demonstrate that deep-learning-based analysis within the CATS framework has the power to reveal synaptic transmission sites, leveraging local context and structural labeling of putative synaptic clefts.

### Synaptic input structure of identified, functionally characterized hippocampal neurons

To combine structural and functional readout, we performed coCATS in organotypic hippocampal slice cultures (**Fig. 3a,b, Supplementary Video 3**) after whole-cell patch clamp recordings. CATS revealed pSCRs and provided comprehensive structural context to individual, electrophysiologically characterized neurons, including DG granule cells, CA1 pyramidal neurons, CA3 interneurons, and CA3 pyramidal neurons, which were filled with fluorophores during recording for later identification (**Fig. 3c-e, Supplementary Fig. 11**). Electrophysiological recordings during and after coCATS label application showed that neuronal activity (induced action potential generation) continued (**Supplementary Fig. 12**), demonstrating that cells were alive and functional at the time of fixation.

**Fig. 3.**
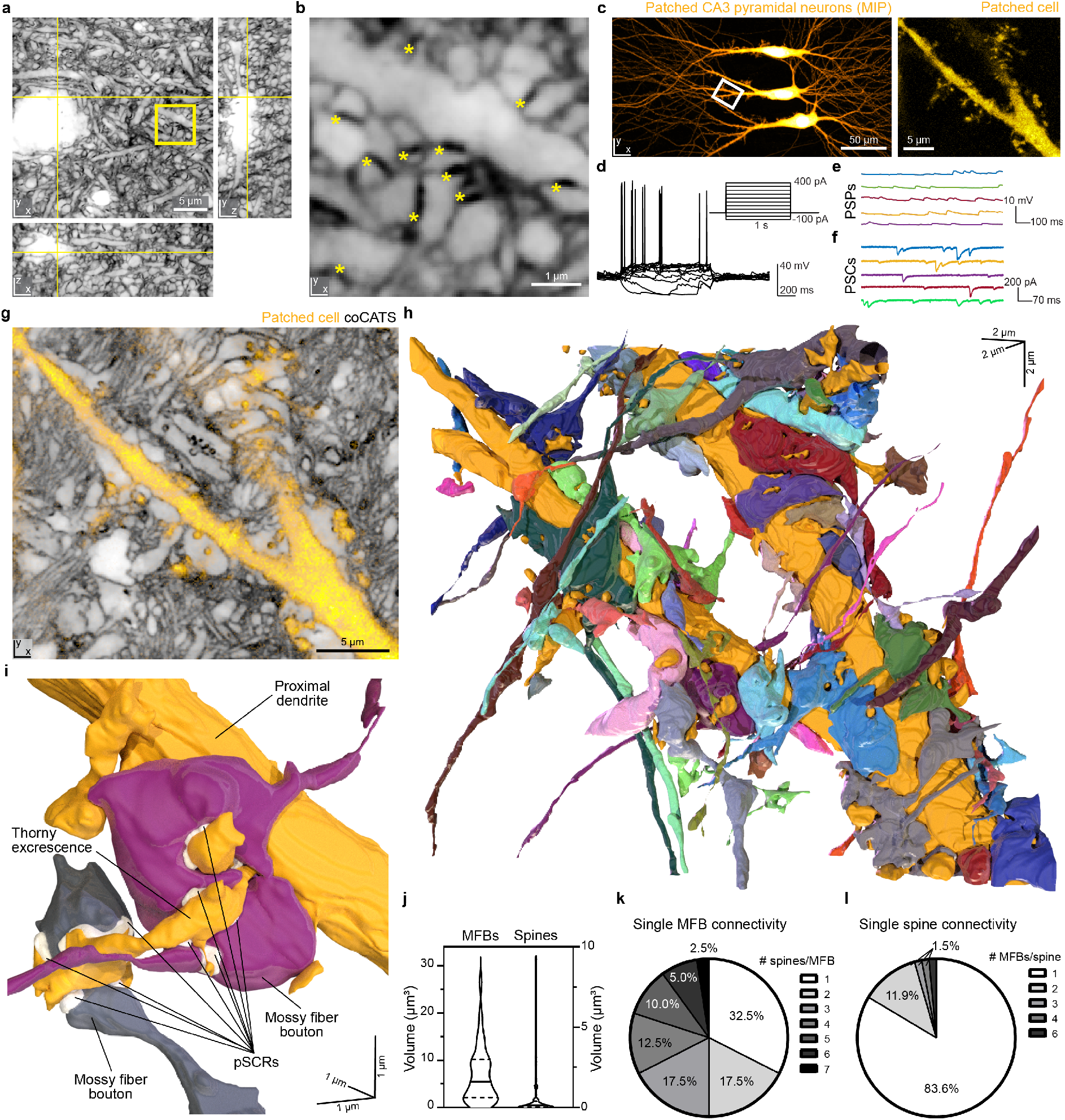
Reconstruction of CA3 pyramidal neuron local input field with coCATS. **a**, Orthogonal views of a coCATS imaging volume recorded with *z*-STED at near-isotropic resolution (N2V, raw data: **Supplementary Fig. 7**) in neuropil of an organotypic hippocampal brain slice. Yellow lines indicate position of displayed planes. **b**, Magnified view of the boxed region in a. Asterisks: pSCRs. **c**, (*Left*) CA3 pyramidal neurons in an organotypic hippocampal slice whole-cell patch-clamped and filled with fluorescent dye (Lucifer yellow). (*Right*) Magnified view of a piece of proximal dendrite in the boxed region. **d**, Action potential response of the middle pyramidal neuron elicited by current injection (inset). **e,f**, Spontaneous post-synaptic potentials (PSPs) and post-synaptic currents (PSCs) recorded from middle pyramidal neuron. **g**, CoCATS (grey, *z*-STED, N2V) overlaid with the intracellular label (yellow, confocal) of the middle pyramidal neuron provides super-resolved information on its local micro-environment. **h**, 3D-rendering of the same proximal dendrite (gold) and 57 structures synaptically connected to it, reconstructed from the volumetric coCATS data. Connectivity was inferred by the presence of pSCRs between the positively labeled dendrite and the respective adjacent structures. **i**, 3D-rendering of two MFBs (violet, grey) forming complex connections with one thorny excrescence of the proximal dendrite. PSCRs are indicated in white. **j**, Violin plots with median (line) and quartiles (dashed lines) of the MFB volumes (n_MFB_=40) contacting the recorded pyramidal neuron and its spines (n_spine_=68). **k,l**, Quantification of connectivity pattern of individual MFBs and pyramidal neuron spines.

CATS visualized electrophysiologically characterized neurons together with surrounding structures at near-isotropic STED resolution, revealing key information missing with sparse positive cellular labeling alone (**Fig. 3c,g**). We set out to determine the complete connectivity structure of a proximal dendrite (**Fig. 3g**). Proximity of pre- and post-synaptic structures is an unreliable predictor of synaptic connectivity^41^. However, with application of deep-learning-based pSCR segmentation followed by manual validation, coCATS allowed us to identify synaptically connected structures (**Supplementary Fig. 13**). We mapped the synaptic input structure of a proximal dendrite in an electrophysiologically characterized CA3 pyramidal neuron (**Fig. 3h, Supplementary Fig. 14**). From the coCATS data, we reconstructed 58 structures (43 MFB- and 14 non-MFB structures synaptically connected to the dendrite of the recorded cell) to clarify the intricate 3D spatial arrangement of individual MFBs and their post-synaptic complex spines (**Fig. 3h,i, Supplementary Video 4**). MFBs contacting the same dendrite displayed a wide range of sizes, with smaller mean volume and larger spread (**Fig. 3j**, 6.85±5.95 µm^3^) than the manually selected MFBs in adult brain in Fig. 2, potentially reflecting an earlier developmental stage^37^ in the ∼20 days *in vitro* slice cultures. The volume distribution of spines (68 reconstructed) on the pyramidal neuron included large spines with highly complex shapes, i.e. quintessential thorny excrescences. However, the size distribution was strongly skewed towards small spines emanating from the shaft, also in contact with MFBs (**Fig. 3j**). We next evaluated connectivity of individual MFBs (**Fig. 3k**). Interestingly, only ∼1/3 of MFBs formed connections with a single spine, whereas synaptic contact with multiple spines was more common, with single boutons connecting to up to 7 different spine structures. Conversely, individual, especially small, spines mostly connected to single MFBs, but some (16.4%), mostly elaborate, complex spines, were synaptically contacted by more than one MFB (e.g. **Fig. 3i)**. We observed a maximum of 6 pre-synaptic boutons for the largest of the post-synaptic structures (**Fig. 3l**). Together, these data shed light on the complex organization of the mossy fiber circuitry, with signal integration at CA3 pyramidal neurons occurring even at the level of individual thorny excrescences. More broadly, it demonstrates the power of CATS to provide quantitative data on structural and functional connectivity.

### Synaptic output structure and differential tissue architecture across regions

We next took advantage of CATS’ contextual information from single synapse to regional scale, characterizing the synaptic output field of an individual DG granule cell in an organotypic hippocampal slice culture. We performed coCATS labeling after electrophysiological recording (**Supplementary Fig. 15**) and biocytin-filling. We followed the main axon, as it travelled from the cell body in the DG granule cell layer through the hilus to the CA3 *stratum lucidum* (**Fig. 4a**). We performed volumetric, isotropically resolving STED imaging around 17 conspicuous pre-synaptic boutons, focusing mostly on complex boutons (**Fig. 4b,c**). While the axon trajectory and bouton structure could be determined already from the super-resolved positive single-cell label, CATS was required to identify post-synaptic partners via pSCR connectivity, evaluated by deep-learning-based segmentation and manual validation, and to reveal structural context (**Fig. 4c, Supplementary Fig. 15,16**). We analyzed complex mossy fiber boutons, but also smaller *en passant* boutons with identified pSCRs. *En passant* boutons displayed a single synaptic transmission site (one pSCR) to thin dendritic structures and lacked filopodia. In contrast, large boutons featured multiple pSCRs and filopodia in both the hilus (4.0±2.0 filopodia per analyzed bouton) and CA3 *stratum lucidum* (8.5±3.4 filopodia per bouton). These structures formed complex synapses with hilar mossy cells and CA3 pyramidal neurons, respectively, identifiable from their cellular morphology and context in CATS staining. We reconstructed synaptic units in the hilus (**Fig. 4d, Supplementary Video 5**) and the CA3 *stratum lucidum* (**Fig. 4e, Supplementary Video 6**), showing the difference in complexity between *en passant* boutons (boutons 2 and 4) and complex boutons (bouton 13). In CA3, we observed connections between bouton 13 and nine neuronal structures (**Fig. 4e**). These included both engulfment of thorny excrescences by the main bouton, and contacts via filopodial extensions. We also observed pSCRs at synapses formed by filopodia, which are thought to mainly contact inhibitory interneurons, remarkably enhancing complexity of the circuitry^42^.

**Fig. 4.**
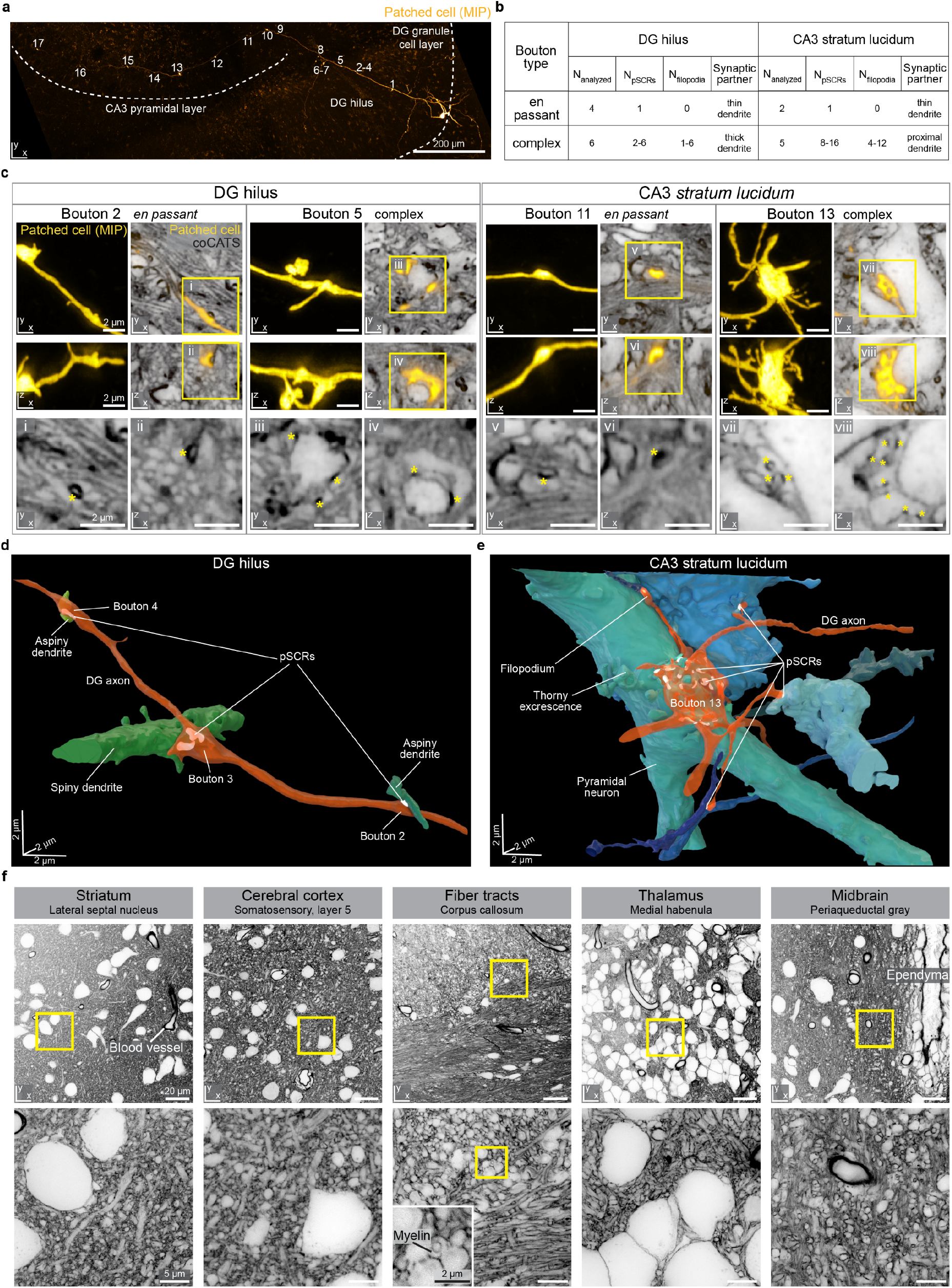
Tissue architecture and single-cell output structure at individual synapse level across brain regions. **a**, Maximum intensity projection (MIP) of a whole-cell patch-clamped and biocytin-filled DG granule cell in organotypic hippocampal slice (confocal). 17 conspicuous boutons are marked along the main axon’s trajectory, projecting as mossy fiber from the DG granule cell layer through the hilus to the CA3 *stratum lucidum*. **b**, Characteristics of analyzed synaptic boutons. **c**, Single *xy*- and *xz*-planes of 4 example super-resolved volumes comprising specific synapses as marked in a, with coCATS (grey, *z*-STED, N2V) revealing local microenvironment of the positively labeled mossy fiber (yellow, *z*-STED, N2V) (raw data: **Supplementary Fig. 7**). (*Bottom*) Magnified views of the coCATS channel with asterisks indicating pSCRs used to identify synaptic partners. **d,e**, 3D-renderings of two axon stretches with boutons, pSCRs, and synaptically connected structures in DG hilus and CA3 *stratum lucidum*. **f**, Architecture of various regions in near-natively preserved brain revealed by coCATS with *in vivo* microinjection. Organization of cell bodies, dendrites, axons, synapses, ependyma around liquor spaces, and blood vessels is visible. Myelinated axons can be distinguished by a fine demarcation of the inner border of the myelin sheath (inset, corpus callosum). (*Top*) Confocal, (*bottom*) *xy*-STED. Raw data.

Seeking to reveal tissue architecture beyond hippocampal circuitry, we returned to *in vivo* coCATS labeling. Microinjection into lateral ventricles or cortex (**Supplementary Fig. 17**) visualized the diversity of cell and tissue architecture in cerebral cortex, different areas of the hippocampus (dentate gyrus, CA1, CA3), striatum, corpus callosum, thalamus, hypothalamus, hindbrain and cerebellum (**Fig. 4f, Supplementary Fig. 18**). Tissue structure was intact beyond ∼200 µm of local damage around the injection site (**Supplementary Fig. 17**). STED disclosed rich structural detail of neuronal and glial processes, synapses, axon bundles, blood vessels, and ependyma in all these regions, with e.g. myelinated axons in the corpus callosum standing out by demarcation of the inner border of myelin sheaths (**Fig. 4f**).

### Large-scale tissue analysis with CATS and expansion microscopy

ExM involves hydrogel embedding, disruption of mechanical cohesiveness of the tissue, and subsequent isotropic swelling, while conserving relative spatial arrangements^5^. This provides effective super-resolution with diffraction-limited readout. It reduces autofluorescence and homogenizes sample refractive index to that of water, clearing the tissue and mitigating aberrations and scattering. Together, this facilitates acquisition of axially extended, super-resolved volumes. We therefore sought to combine CATS’ capability to decode tissue architecture with the strengths of ExM. Expansion requires a label that is faithfully retained in the hydrogel and is minimally affected by the radical chemistry during gel polymerization and heat/chemical denaturation during preparation. Biotin fulfills these criteria, such that we screened for biotin-containing coCATS labels (**Supplementary Fig. 1**). We found that an additional chemical group was required to ensure sufficient extra-to-intracellular contrast and chose PEG12. We live labeled organotypic hippocampal slice cultures with NHS-PEG12-biotin, used heat/chemical denaturation (**Fig. 5a,b, Supplementary Fig. 19**) or enzymatic digestion (**Supplementary Fig. 19**) to disrupt tissue cohesiveness. We expanded ∼4-fold with the magnified analysis of proteomes (MAP)^6^ and protein-retention ExM^8^ approaches, respectively, followed by application of fluorophore-conjugated streptavidin to visualize the extracellular label. This provided signal amplification and flexibility with downstream processing. We recorded confocal stacks of ∼400 µm axial range, showing that it is straightforward to obtain super-resolved context over 100 µm depth at native tissue scale.

**Fig. 5.**
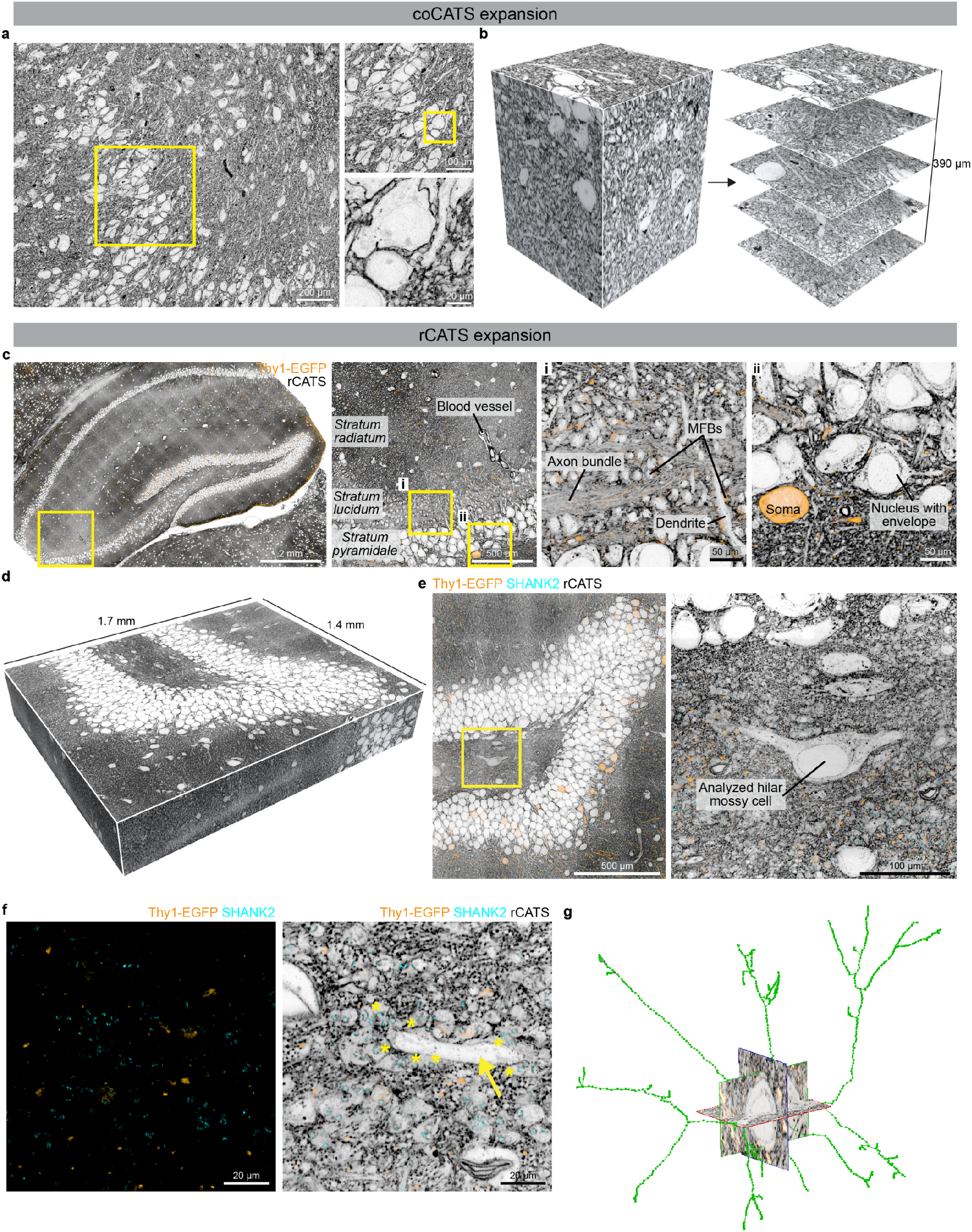
Large-scale imaging of tissue context with expansion microscopy. **a**, Organotypic hippocampal brain slice live labeled with coCATS (NHS-PEG_12_-biotin probe) and ∼4-fold expanded with MAP^6^. Confocal image with two progressive zoom-ins (raw data). Scale bars refer to tissue size after expansion throughout. **b**, Extended depth imaging in ∼4-fold expanded organotypic hippocampal brain slice after live coCATS labeling, showing the imaging volume (*left*) and five single planes at progressively larger depths (*right*). Axial imaging range in confocal readout (N2V) was ∼400 µm, corresponding to ∼100 µm in the original tissue. **c**, Brain tissue section from previously perfusion-fixed Thy1-EGFP+ adult mouse with sparse neuronal labeling by cytosolic EGFP expression (visualized with immunostaining, orange), labeled with rCATS and ∼4-fold expanded by protein-retention ExM^8^, showing full hippocampal area. Zoomed views in the CA3 at two different magnifications show that rCATS delineates tissue context from gross organization down to sub-cellular morphology (confocal, raw data). Zoom i shows mossy fiber boutons as globular structures among axon (mossy fiber) bundles in the *stratum lucidum*, zoom ii shows the arrangement of cell bodies and neuropil at the outer border of the *stratum pyramidale*. **d**, 3D-representation of a volume of the DG crest of a previously perfusion-fixed Thy1-EGFP+ adult mouse. After rCATS labeling (grey, N2V), immunostaining for SHANK2 (cyan, N2V) and EGFP (orange, N2V), the sample was 4.5-fold expanded and imaged with high-speed spinning disc microscopy. The displayed volume corresponds to 303×371×70 µm^3^ in original tissue volume. **e**, Single *xy*-plane of the data represented in d with zoom-in on the soma of a hilar mossy cell. **f**, Different plane from the same volume showing immunostainings alone and overlaid with rCATS at higher magnification. The yellow arrow indicates a dendrite belonging to the mossy cell displayed in e, lined by MFBs with SHANK2 located at the synaptic transmission sites. Yellow asterisks highlight a subset of MFBs in contact with the mossy cell dendrite. **g**, Skeletonization of the major branches of the hilar mossy cell in e,f from rCATS data.

For several important preparations, in particular previously fixed brain, it is not possible to perform extracellular labeling while the tissue is alive. We therefore screened binders to ECS-resident molecules that were widely and homogeneously distributed in mouse brain (resident CATS, rCATS). Different polysaccharide-binding proteins showed distinct labeling patterns, highlighting the molecular diversity in the ECS (**Supplementary Fig. 20**). Wheat germ agglutinin (WGA) binds to N-acetyl-D-glucosamine and sialic acid and has been used in different organs to outline blood vessels or cell bodies^43,44^. Labeling fixed mouse brain tissue with fluorescently marked WGA revealed hippocampal architecture clearly (**Fig. 1b**). Myelinated axons were distinguishable in STED mode, as validated by specific staining (**Supplementary Fig. 21**), as well as carbohydrate-rich nuclear pores. However, WGA features few lysines for hydrogel anchoring, resulting in poor retention upon expansion (**Supplementary Fig. 22**). To make rCATS compatible with ExM, we developed a dedicated signal retention strategy (**Supplementary Fig. 22**), transferring information from biotinylated WGA to acrylamide-modified streptavidin copolymerizing with the gel. Large-scale readout of expanded samples with spinning disc confocal microscopy allowed high resolution visualization of tissue architecture (**Fig. 5c, Supplementary Fig. 23**). To illustrate the rich information contained in this type of data, we imaged a 1.4×1.7×0.32 mm^3^ (post-expansion; expansion factor 4.5; 303×371×70 µm^3^ pre-expansion; ∼1 TB) volume of the DG crest and enclosed hilus, wherein rCATS provided structural context to sparse Thy1-EGFP labeled neurons and excitatory synapses labeled for SHANK2 (**Fig. 5d-f**). Given the large scale of the data, we skeletonized the major dendritic arborizations of an unlabeled example neuron from the rCATS signal. This cell, identified as a mossy cell by its morphology and the presence of multiple MFBs in contact with its dendrites, can be studied amidst its 3D-context, demonstrating the utility of rCATS for unbiased imaging and analysis of any neuronal population in the tissue (**Fig. 5f,g**).

### CATS in human brain tissue

Analysis of human clinical brain samples largely relies on conventional histology stainings, such as hematoxylin and eosin, that coarsely represent tissue architecture. To test whether CATS can be adopted to human samples, we obtained fixed hippocampal tissue extracted from a 36 year-old male patient undergoing epilepsy surgery. We applied rCATS at confocal resolution together with immunolabeling of neuronal processes (microtubule-associated protein 2, MAP2) and excitatory synapses (HOMER1). Also in the human samples, rCATS labeling revealed contextual information and differential architecture in the layers of the DG (**Fig. 6a,b**). In addition, comprehensive visualization by rCATS allowed detailed, yet straightforward, assessment of tissue preservation, the major quality determinant for microanatomical studies of clinical brain material. In contrast, immunostainings alone without rCATS made it challenging to distinguish effects of tissue degradation due to the sparse distribution patterns of target molecules. RCATS is thus a valuable resource for studying tissue structure and single-cell morphology in clinical specimens of healthy and diseased individuals.

**Fig. 6.**
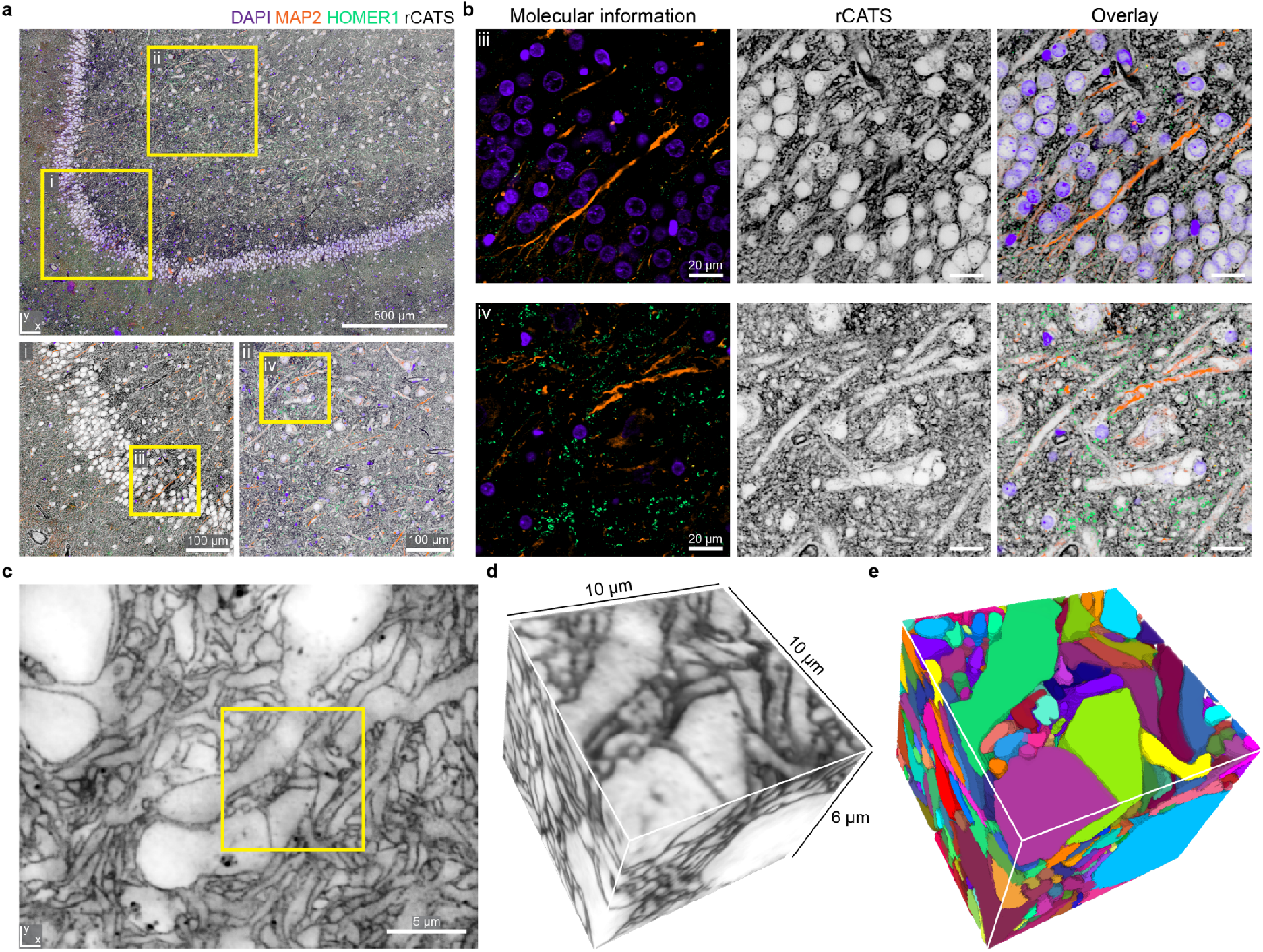
Tissue architecture in human nervous tissue. **a**, RCATS (grey) in the DG-region of a human hippocampal surgery specimen with additional staining for dendrites (MAP2, orange), excitatory synapses (HOMER1, green) and nuclei (DAPI, purple). (*Top*) Confocal, with magnified views of boxed regions (*bottom*). **b**, Magnified views of the boxed regions iii and iv in a. (*Left*) Molecular stainings alone, (*middle*) tissue architecture revealed by rCATS, (*right*) overlay. Raw data. **c**, CoCATS in human cerebral organoid. Single plane of super-resolved volume (*z*-STED, N2V, adaptive histogram equalization). **d**, Subvolume of the same dataset, as indicated in c. **e**, Dense tissue reconstruction with coCATS via manual segmentation of the volume in d.

Finally, we sought to demonstrate the applicability of CATS to human cerebral organoids. Organoids are emerging as an experimentally tractable human system for studying brain development and disease mechanisms^45^. We asked whether CATS could be extended to densely reconstruct the cellular constituents of an organoid volume. We chose coCATS, as it is independent of the deposition of extracellular matrix molecules in this early development model system. Using STED at near-isotropic resolution allowed dense cellular segmentation (**Fig. 6c-e, Supplementary Video 7**), making CATS useful for cell and tissue phenotyping also in this sample type. The organoid showed lower complexity than the other sample types analyzed. However, this proof-of-principle experiment paves the way towards large-scale dense reconstruction of complex tissue samples with light microscopy.

## Discussion

Here we developed CATS, a labeling, imaging, and analysis platform to map brain tissue architecture across spatial scales with light microscopy. CATS places cells and specific molecules into their tissue context, and allows quantifying neuronal connectivity and reconstructing tissue structure down to subcellular nano-morphology, including single synapses. Designed for fixed rather than living tissues, it facilitates analysis of diverse specimens and extended volumes, using readily available super-resolution approaches, specifically STED and expansion microscopy. Contrary to the selective representation with positive cellular labeling, CATS displays the tissue with all its cellular structures in an unbiased fashion. CATS labels molecules in extracellular space and on cell surfaces, with the structural imaging channel remaining free from intracellular complexity. This creates a clear boundary between cells and allows distinguishing cellular structures at high contrast in extremely dense brain tissue, even when read out at diffraction-limited resolution or comparatively moderate resolution increase over the diffraction limit. Together with broad compatibility with standard molecular labeling techniques, this property will facilitate widespread incorporation into tissue analysis workflows, dramatically advancing their information content.

We employed two labeling strategies, coCATS and rCATS, to cater for diverse brain tissue preparations, including native rodent brain, organotypic slice cultures, previously fixed mouse and human brain tissue, and human cerebral organoids. Broad labeling of extracellular molecules in coCATS and the resulting high labeling density enabled reconstruction of single synapse morphology, whereas rCATS extended usefulness to sample types that do not allow live labeling. CATS is a technologically straightforward approach to 3D tissue reconstruction in the vast number of applications where EM resolution is not essential and adds super-resolved molecular information to 3D reconstruction. By directly bridging spatial scales (mm–nm), it avoids complex correlative workflows that require reconciling different sample preparation and imaging modalities.

Obvious improvements include specifically engineering labels for enhanced hydrogel retention^46^ or signal amplification and, in rCATS, multi-component labeling of extracellular molecules, as well as increasing readout speed with light-sheet microscopy^14^. CATS paves the way towards development of molecularly informed, light-microscopy based connectomics. For this, specifically increasing optical resolution or expansion factors^47–50,29,28^ should allow tracing also the thinnest of neuronal structures, finer than the resolution employed here.

We used the hippocampal mossy fiber circuitry as first application target. Quantifications of single-bouton geometry and connectivity are in line with benchmark EM data^36,38,39^. However, in contrast to EM, CATS enabled straightforward incorporation of specific molecular information in 3D-reconstructions and massively reduced requirements in time, personnel and equipment over classical serial-sectioning EM studies. For example, imaging the three volumes used to reconstruct the 30 MFBs in **Fig. 2** required only ∼4h hands-on sample preparation and 3×1.5 h total imaging time for the 3-channel measurement.

Taking advantage of comprehensive tissue visualization, we addressed a long-standing challenge in brain tissue imaging by applying CATS to decode synaptic connectivity. A surprising, but powerful discovery is that coCATS unveils putative synaptic cleft regions (pSCRs) by a prominent labeling pattern within its structural context. These are detectable with specifically tailored machine learning analysis even in absence of molecular staining. Accordingly, pSCRs can be leveraged to infer and quantify synaptic connections, and identify synaptic partners among neighboring structures. We used this to reconstruct the local synaptic input structure of an identified CA3 pyramidal neuron and to characterize the synaptic output structure of a DG granule cell when following its main axon across the hippocampus. In both cases our analysis revealed stunning complexity, showing CATS’ power to unravel the structural correlates of diverging and converging signal integration in the central nervous system.

Our analysis presents one of the largest datasets of local mossy fiber connectivity reconstruction to date. Throughput of 3D reconstruction was limited by manual volume segmentation. Overall throughput will substantially benefit from replacing manual cell shape segmentation with deep-learning-based approaches adopted from EM connectomics^51–53^, as already employed in super-resolution reconstruction of living brain tissue^24^. This will make large-scale studies of local connectivity, complete neuronal synaptic input or output fields, and neuron-glia interplay^21^ feasible. We furthermore expect CATS to seamlessly integrate with complementary technologies, such as calcium imaging or viral circuit tracing^54,55^, similar to the combined structural and functional characterization demonstrated here with whole-cell patch clamp recordings. CATS will be an important tool to clarify how tissue architecture and synaptic connectivity are remodeled in response to neuronal activity, during development, or in neurodevelopmental or neurodegenerative disease models. CATS furthermore opens the door to unbiased phenotyping of cell and tissue structure of rodent and patient-derived human samples, shedding new light on tissue architecture, cell-cell interactions, and subcellular morphology both in healthy and diseased brain. High throughput, easy adoptability, and seamless pairing of structural data with molecular and functional information puts CATS in an excellent position to clarify structure-function or genotype-to-tissue-phenotype relationships. Taken together, CATS is a powerful tool for phenotyping brain and provides unprecedented views into cellular microenvironments both in health and disease.

## Acknowledgements

We thank Jakob Vorlaufer, Nathalie Agudelo-Dueñas, Wiebke Jahr and Andreas Wartak for microscope maintenance and troubleshooting, as well as Caroline Kreuzinger and Anna Freeman for technical assistance. We gratefully acknowledge Eder Miguel for setting up webKnossos and Marek Šuplata for computational support and hardware control. We are grateful to Ryuichi Shigemoto and Bernd Bickel for generous support, and Michael Sixt and Scott Boyd (Stanford University) for discussions and critical reading of the manuscript. PSD95-HaloTag mice were kindly provided by Seth Grant (University of Edinburgh). We acknowledge expert support by the scientific service units of the Institute of Science and Technology Austria, including scientific computing, imaging and optics, preclinical, and life science facilities, and by the Miba machine shop.

We gratefully acknowledge funding by the following sources:

Austrian Science Fund (FWF) grant I3600-B27 (JGD)

Austrian Science Fund (FWF) grant DK W1232 (JGD)

Austrian Science Fund (FWF) grant Z 312-B27, Wittgenstein award (PJ)

Gesellschaft für Forschungsförderung NÖ (NFB) grant LSC18-022 (JGD)

European Union’s Horizon 2020 research and innovation programme, European Research Council (ERC) grant 715508 – REVERSEAUTISM (GN)

European Union’s Horizon 2020 research and innovation programme, European Research Council (ERC) grant 692692 – GIANTSYN (PJ)

Marie Skłodowska-Curie Actions Individual Fellowship 101026635 under the EU Horizon 2020 program (JFW)

## Author contributions

JMM and JGD designed the study, experiments, and analysis and interpreted data. JMM performed experiments, analysis, proofreading, and visualization, prepared figures and contributed to manuscript writing. JL designed and performed analysis, visualized and interpreted data, prepared figures and contributed to manuscript writing. PV supported experiments. HK performed stereotactic injections. JFW performed patch-clamp experiments. AC performed manual segmentations. CS supported image analysis. AV, KR, TC, and SS provided human brain surgery specimens. GN advised on and provided human cerebral organoids. PJ supervised patch clamp experiments and advised on synaptic neuroscience and hippocampal circuitry. JGD initiated and supervised the study. JGD wrote the manuscript with critical input from all authors.

