## Supplementary Information for "Uncovering brain tissue architecture across scales with super-resolution light microscopy"

Julia M. Michalska[1], Julia Lyudchik[1], Philipp Velicky[1], Hana Korinkova[1], Jake F. Watson[1], Alban Cenameri[1], Christoph Sommer[1], Alessandro Venturino[1], Karl Roessler[2], Thomas Czech[2], Sandra Siegert[1], Gaia Novarino[1], Peter Jonas[1], Johann G. Danzl[1]

[1]Institute of Science and Technology Austria, Am Campus 1, 3400 Klosterneuburg, Austria

[2]Medical University of Vienna, Department of Neurosurgery, Waehringer Guertel 18-20, 1090 Vienna, Austria

**This pdf file contains:**

Supplementary Figures 1-23 plus captions

Captions for Supplementary Videos 1-7

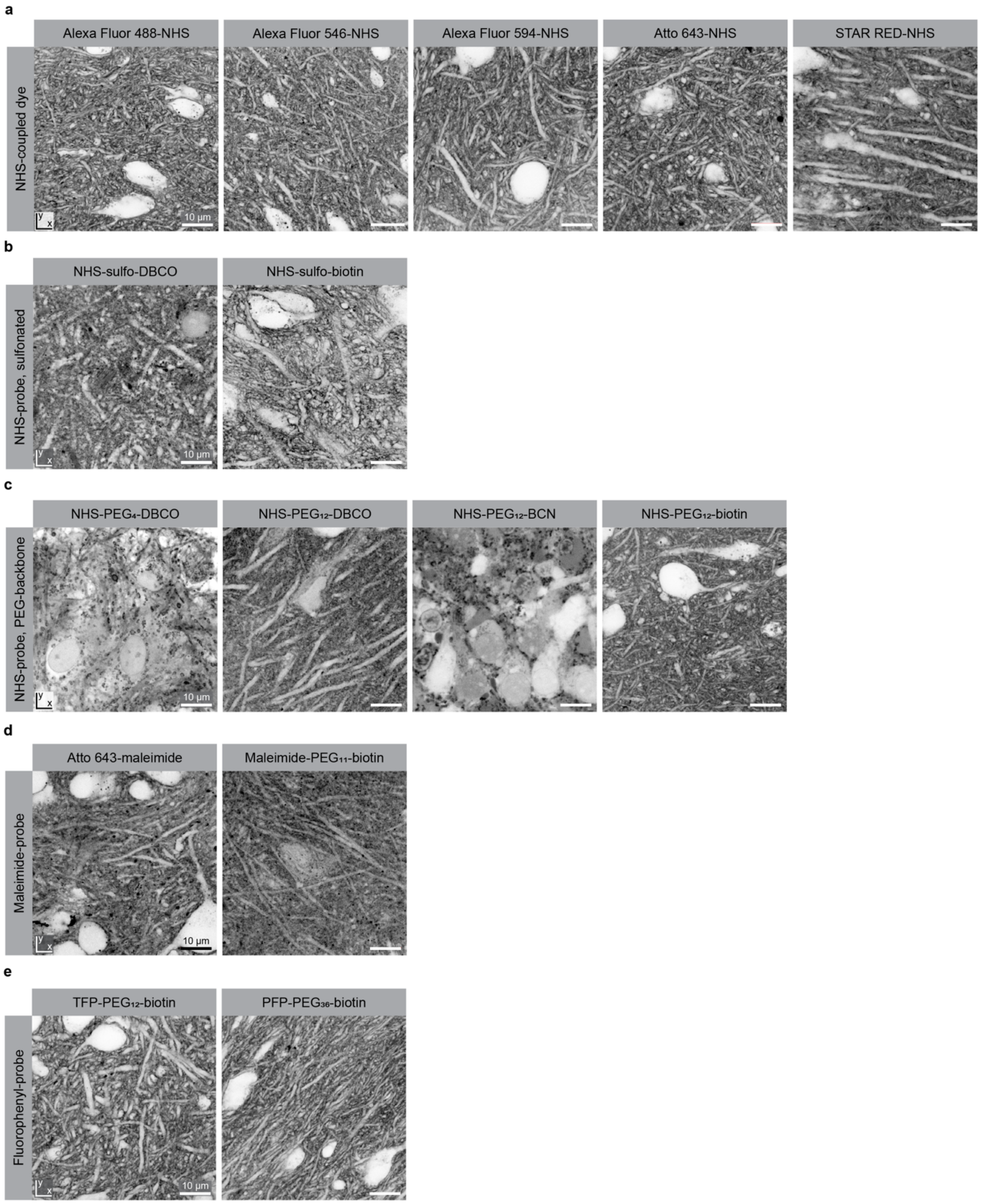


**Supplementary Fig. 1| Screening for coCATS labeling probes.** *Rationale:* Screening was performed in organotypic hippocampal brain slices (15-30 DIV) to identify compounds for coCATS that are i) compatible with live labeling, ii) provide high extra- to intracellular contrast, iii) mediate covalent attachment to molecules in the extracellular space and on cell surfaces, iv) have sufficient tissue penetration, v) achieve homogeneous/high-density labeling and vi) are compatible with downstream super-resolution imaging, specifically STED and expansion microscopy. For covalent attachment, amine-reactive groups (N-Hydroxysuccinimidine (NHS), tetrafluorophenyl (TFP), and perfluorophenyl

(PFP)) and sulfhydryl-reactive maleimide were tested. Readout labels comprised fluorescent dyes or moieties for downstream targeting with fluorescent readout probes (biotin, click chemistry). Directly dye labeled samples were imaged without permeabilization to avoid additional processing steps that may affect structural preservation. For biotin and click readout, the sample was permeabilized. Biotin readout was particularly useful for experiments involving expansion microscopy, as post-expansion fluorophore addition avoids damage to fluorophores by hydrogel radical chemistry or harsh denaturation steps and provides signal amplification. Samples were live incubated with coCATS labels, followed by immersion-fixation, and imaging in confocal mode. **a,** Hydrophilic, negatively charged, NHS-coupled dyes yield high extra- to intracellular contrast and homogeneous labeling. **b,** Sulfo-groups increase hydrophilicity and extra- to intracellular contrast for readout moieties that do not provide membrane impermeability by themselves. Dibenzocyclooctin (DBCO) is a reactive cycloalkane for click-chemistry mediated readout. Its NHS ester derivative did not produce satisfactory results, as DBCO is lipophilic and led to aggregate formation in tissue. In contrast, biotin, with subsequent readout via streptavidin, proved to be an excellent readout moiety. **c,** A polyethyleneglycol (PEG) backbone similarly increases hydrophilicity. We observed that a chain length of 11-12 PEG molecules, but not 4 PEG molecules, was sufficient for mediating high extra- to intracellular contrast. Biotin was preferable over DBCO or another click-chemistry compound, bicyclononyne (BCN), also in this case, as it did not produce aggregates in the tissue. **d,** Maleimide compounds covalently react with sulfhydryl groups. The compounds tested here yielded pronounced extracellular labeling but produced a less homogeneous and more granular staining pattern compared to amine reactive compounds. **e,** TFP and PFP are amine reactive compounds that are more resistant to hydrolysis than NHS esters. They produce high quality labeling. The long PEG chain in PFP-$PEG_{36}$-biotin was not necessary for ensuring extra- to intracellular contrast and merely increased probe size, potentially hampering tissue penetration. All scale bars: 10 μm.

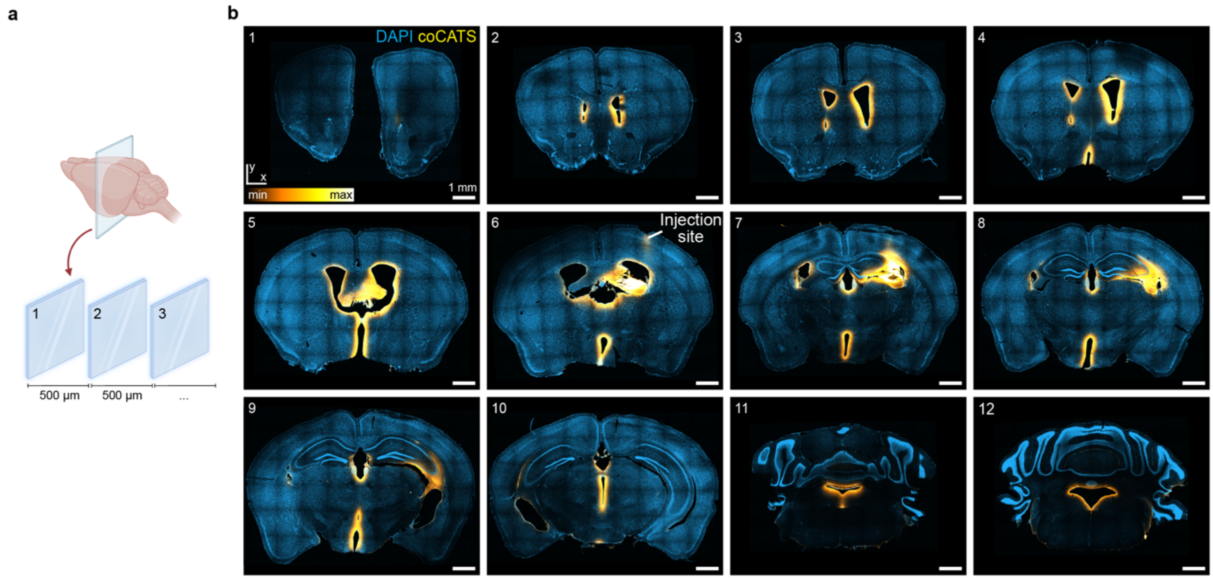


**Supplementary Fig. 2| CoCATS labeling pattern after *in vivo* microinjection into the lateral ventricle.** CoCATS labeling solution was delivered by *in vivo* microinjection into the left lateral ventricle of an adult anesthetized mouse, followed by a 40 min incubation and perfusion-fixation. **a,** Schematic of serial sectioning scheme of the brain after perfusion-fixation. The entire brain was sectioned into 50 µm thick coronal slices. Every 10$^{th}$ slice, spaced 500 µm apart, was used for overview imaging in panel b. **b,** Tile scans of coronal sections, counter-stained with DAPI (blue), acquired with a spinning disc confocal microscope. Intensity lookup table for CATS (yellow) is not inverted. (1) No staining is visible in the anterior sections. (2,3) Labeling commences in both hemispheres around the anterior region of the lateral ventricle where parts of the caudoputamen and lateral septal nucleus line the lateral ventricle. (4) By diffusion into the third ventricle, tissue adjacent to it, including in the hypothalamus, is labeled. (5) Further labeling of parts of the striatum, thalamus and hypothalamus, as well as fiber tracts, is visible around the lateral and third ventricles. (6-9) The injection site is visible close to the cortical surface. Strong labeling of the CA3 can be seen throughout the majority of the hippocampus with decreasing intensity towards the ventral region. (10) Labeling of tissue around the third ventricle, including dorsal hypothalamus. (11,12) Diffusion into fourth ventricle leads to staining of parts of the medulla, pons, and a portion of the central lobule of the cerebellar cortex.

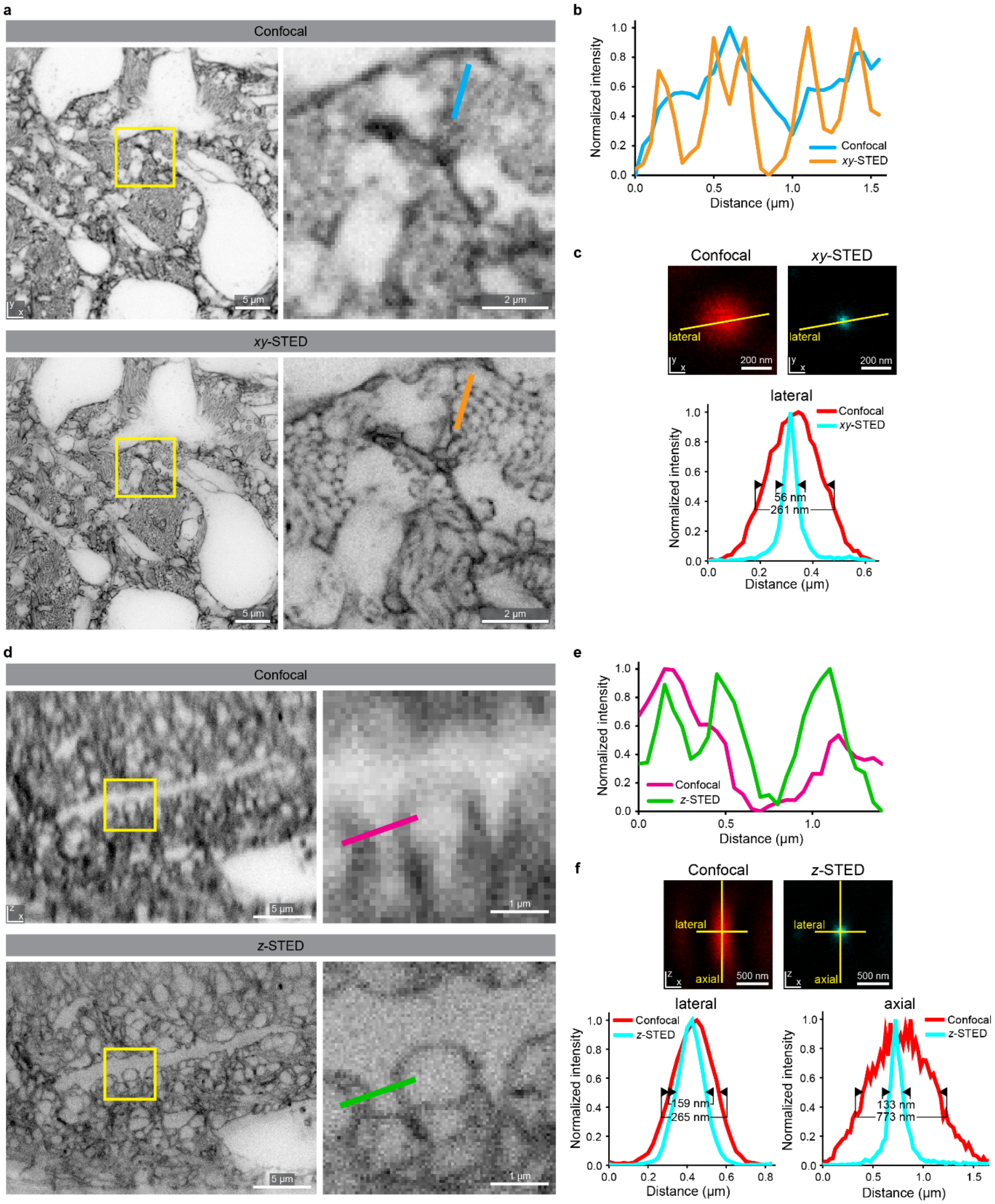


**Supplementary Fig. 3| Improved visualization of tissue architecture with STED microscopy in coCATS. a,** CoCATS in adult mouse hippocampus (DG hilus) with labeling by *in vivo* microinjection into the lateral ventricle. Overview and magnified view of the same region in confocal (*top*) and *xy*-STED mode (*bottom*) with lateral resolution increase. Tissue structure is visible more clearly in STED mode. For example, individual axons are discernible as fine rings only in STED mode. **b,** Line profile (width=3 pixels) in a region of axon bundles as indicated by the line in a, revealing individual axons in STED but not in confocal mode. **c,** Confocal and STED images of a single 40 nm Crimson bead with line profiles and full width at half maximum (FWHM). STED power was the same as in panel a. **d,**

Single axial sections in the neuropil of an organotypic hippocampal slice. Same region imaged in confocal (*top*) and *z*-STED mode (*bottom*). Resolution increase is stronger in the axial direction, yielding near-isotropic resolution with overview and magnified view of the boxed region. STED performance is high in the central ~10-15 µm of the axial range, for which correction of spherical aberrations was set by the objective's correction collar. Decreasing STED performance above and below reflects the well-known sensitivity of the *z*-STED pattern to spherical aberration. **e,** Line profile (width=3 pixels) as indicated in c. **f,** Confocal and STED images of a single 40 nm Crimson bead with corresponding line profiles. STED power was the same as in panel d. All data acquired with the same high-numerical aperture (NA=1.35) silicone oil immersion objective.

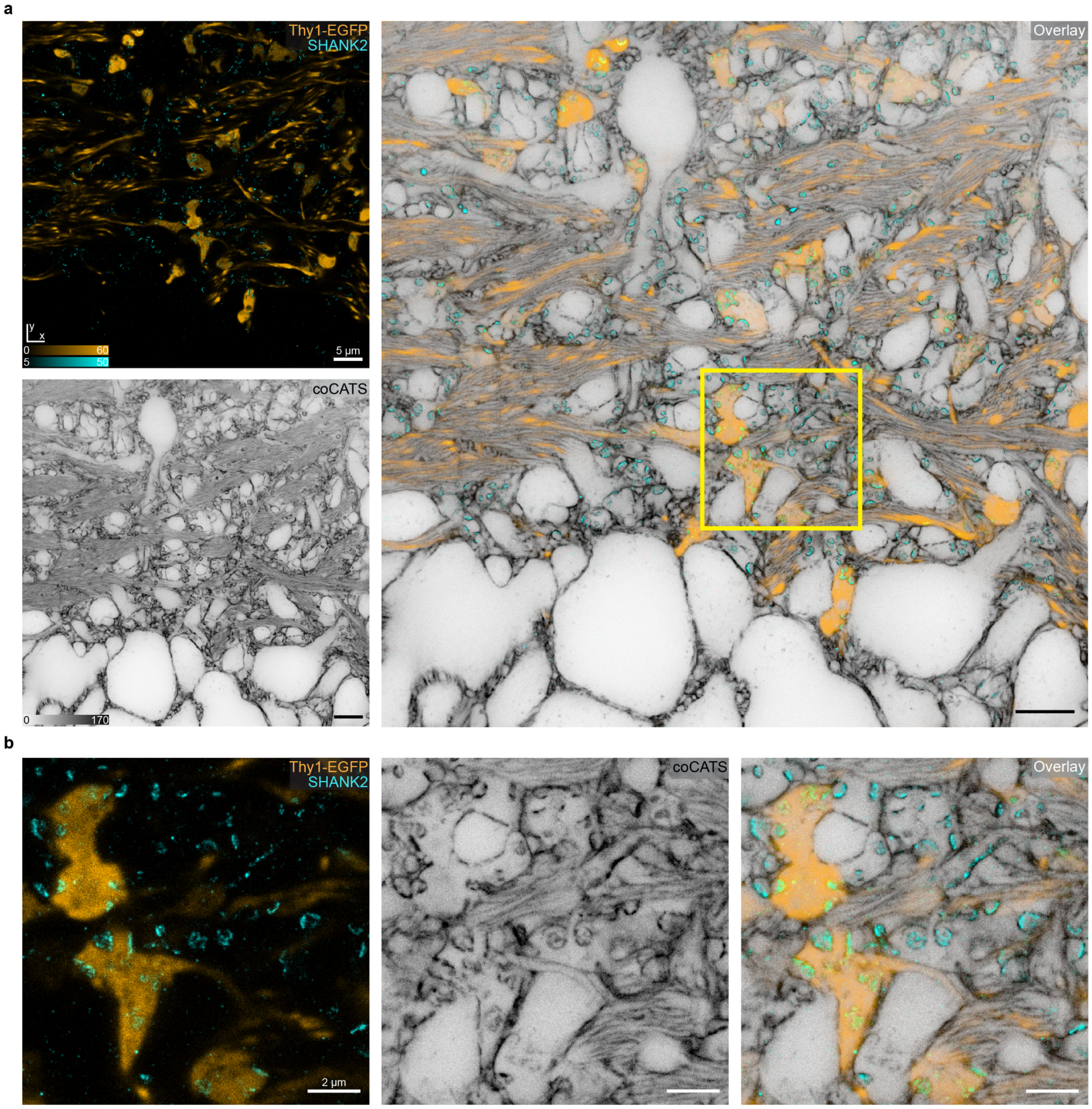


**Supplementary Fig. 4| Information gain with CATS over sparse neuronal labeling. a,** CoCATS (grey, *xy*-STED) labeling by *in vivo* microinjection into the lateral ventricle of an adult mouse combined with a sparse genetic marker (Thy1-EGFP, yellow, confocal, immunostaining for EGFP) and a synaptic marker (immunostaining for SHANK2, *xy*-STED) shows the gain in information provided by CATS. **b**, Magnified view of the yellow box in a, showing mossy fiber boutons, some of which are positively labeled via EGFP expression. When using the sparse genetic marker alone, many synapses (indicated by presence of SHANK2) cannot be assigned within the tissue's structure. CATS, in contrast, reveals not only the positively labeled, but all cellular structures.

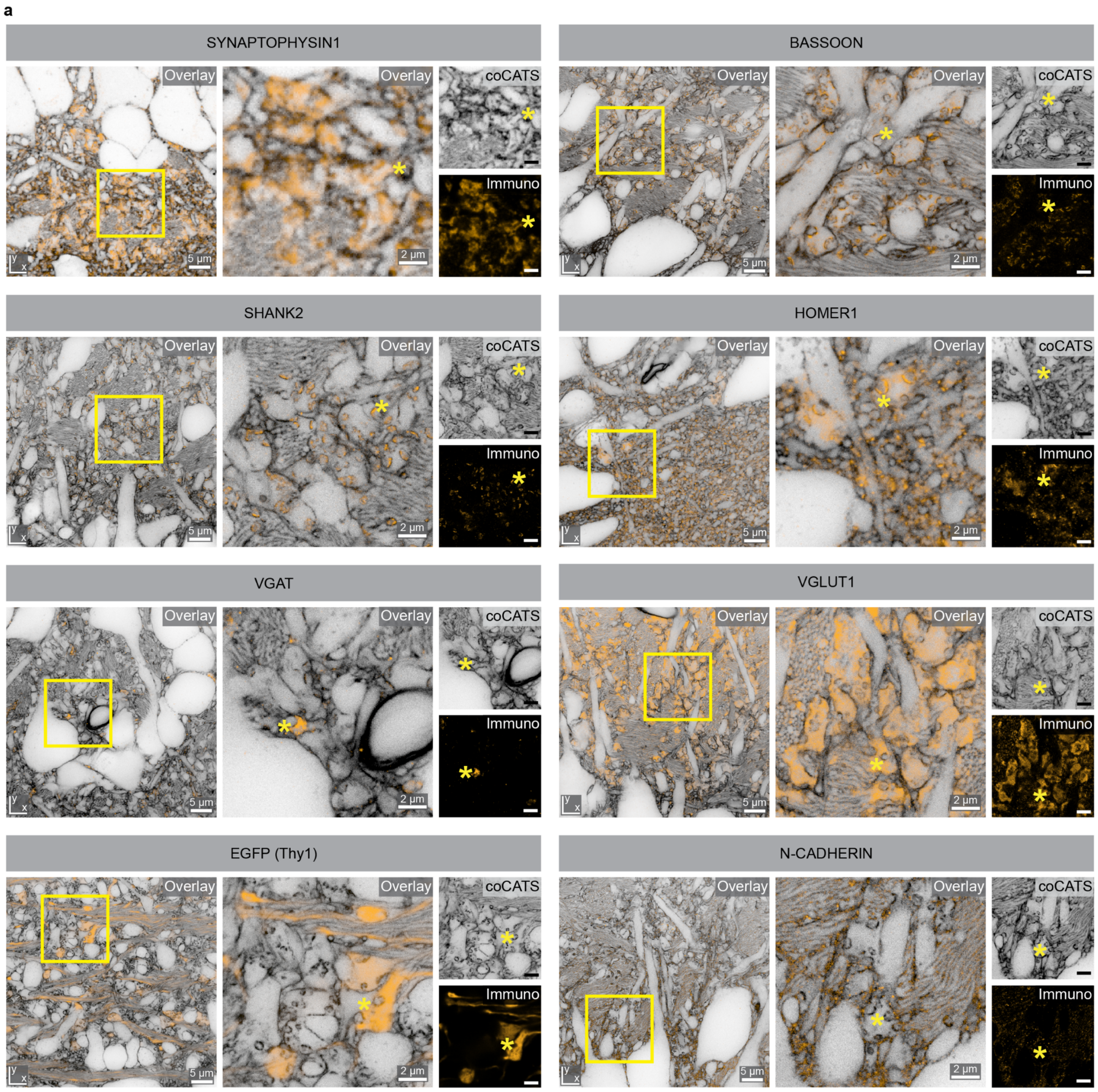


**Supplementary Fig. 5| Validation of putative synaptic cleft region (pSCR) location by immunostaining.** CoCATS in hippocampus labeled via *in vivo* microinjection into adult mouse brain, followed by transcardial fixative perfusion. Immunostaining for the pre-synaptic markers BASSOON (scaffolding protein in active zone), SYNAPTOPHYSIN (synaptic vesicles), VGLUT1 (vesicular glutamate transporter, excitatory synapses), VGAT (vesicular GABA transporter, inhibitory synapses), and the post-synaptic markers SHANK2 (post-synaptic density protein, excitatory synapses) and HOMER1 (post-synaptic density protein, excitatory synapses) reveal their spatial relationship with pSCRs (asterisks). The fact that pSCRs are present both in proximity to VGLUT1 and VGAT indicates that pSCRs can be detected both in excitatory and inhibitory synapses, respectively, with coCATS. Sparse cytosolic EGFP expression in dentate gyrus granule cells (Thy1-EGFP+ mouse line, EGFP detected with immunostaining) highlights a subset of mossy fiber boutons, validating the location of pSCRs with respect to MFBs in this sparse subset. Immunostaining for N-CADHERIN, a marker for cell-cell adhesions, showed little co-localization with pSCRs, indicating that *puncta adherentia* are largely distinct from pSCRs observed with coCATS labeling. Data were acquired with a STED microscope using an *xy*-STED pattern for lateral resolution increase, except for the SYNAPTOPHYSIN1 panel (immunostaining, coCATS label: confocal), and the EGFP signal in the Thy1-EGFP panel (EGFP: confocal; coCATS: *xy*-STED).

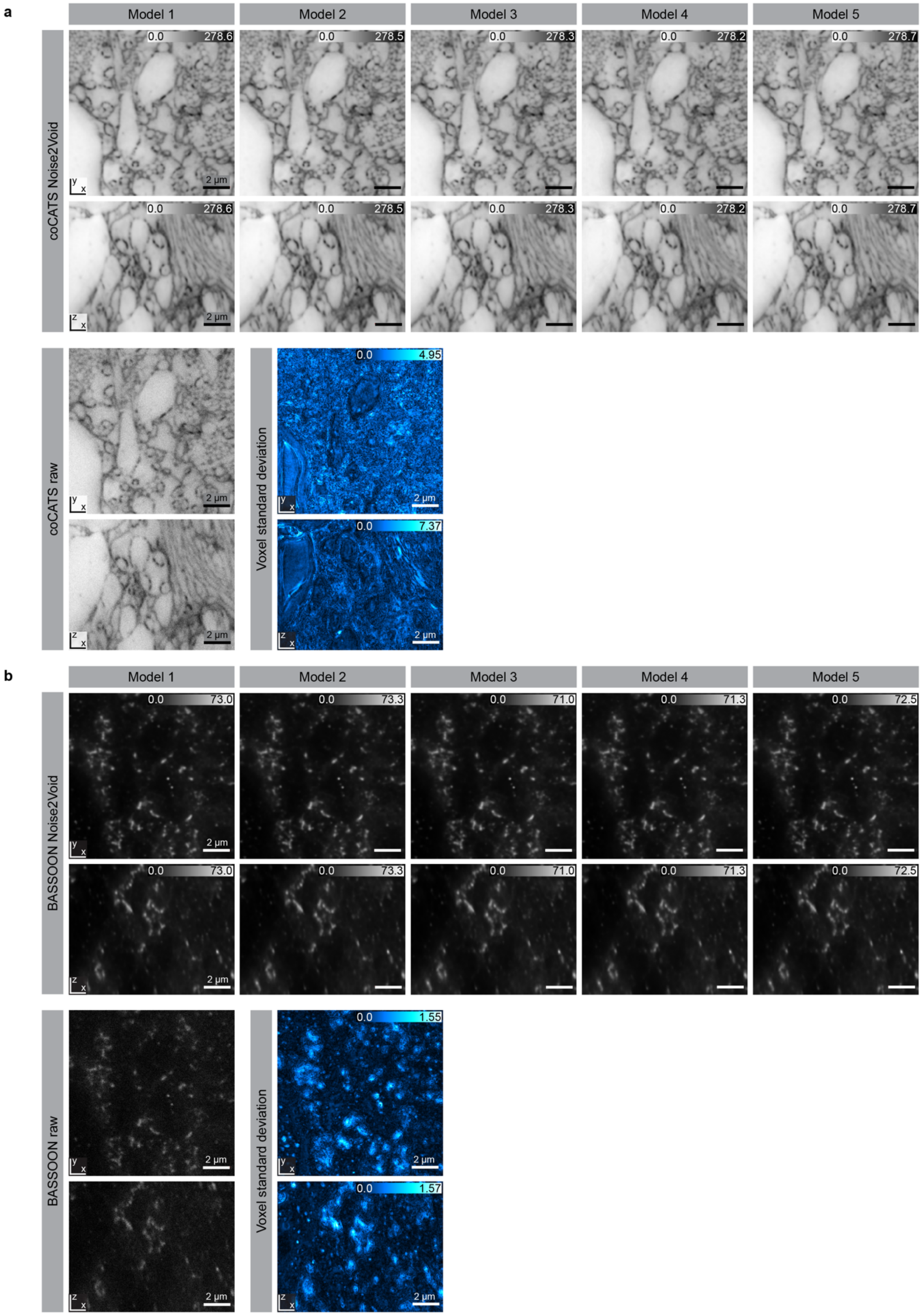


**Supplementary Fig. 6| Validation of Noise2Void (N2V) network prediction for denoising CATS and immunostaining data.** Single *xy*- and *xz*-planes of **a,** coCATS labeling and **b,** super-resolved BASSOON immunostaining after *in vivo* microinjection and perfusion-fixation of an adult mouse.

Volumetric imaging data was acquired with a STED microscope, using *z*-STED pattern for resolution improvement. Data denoised with 5 independent N2V network models, raw data, and voxel-wise standard deviation of the 5 independent N2V models are shown. The voxel-wise standard deviation reflects the standard deviation from the mean of intensity at every voxel across the 5 N2V network models. Highest disagreement is seen in the coCATS channel for areas with low intensity, such as cell bodies, and for BASSOON at the border of immunostaining signals. Note the high degree of similarity of the different N2V outcomes both for the CATS and the immunostaining channel, also reflected by the overall low standard deviation values between the models. Scale bars: 2 μm.

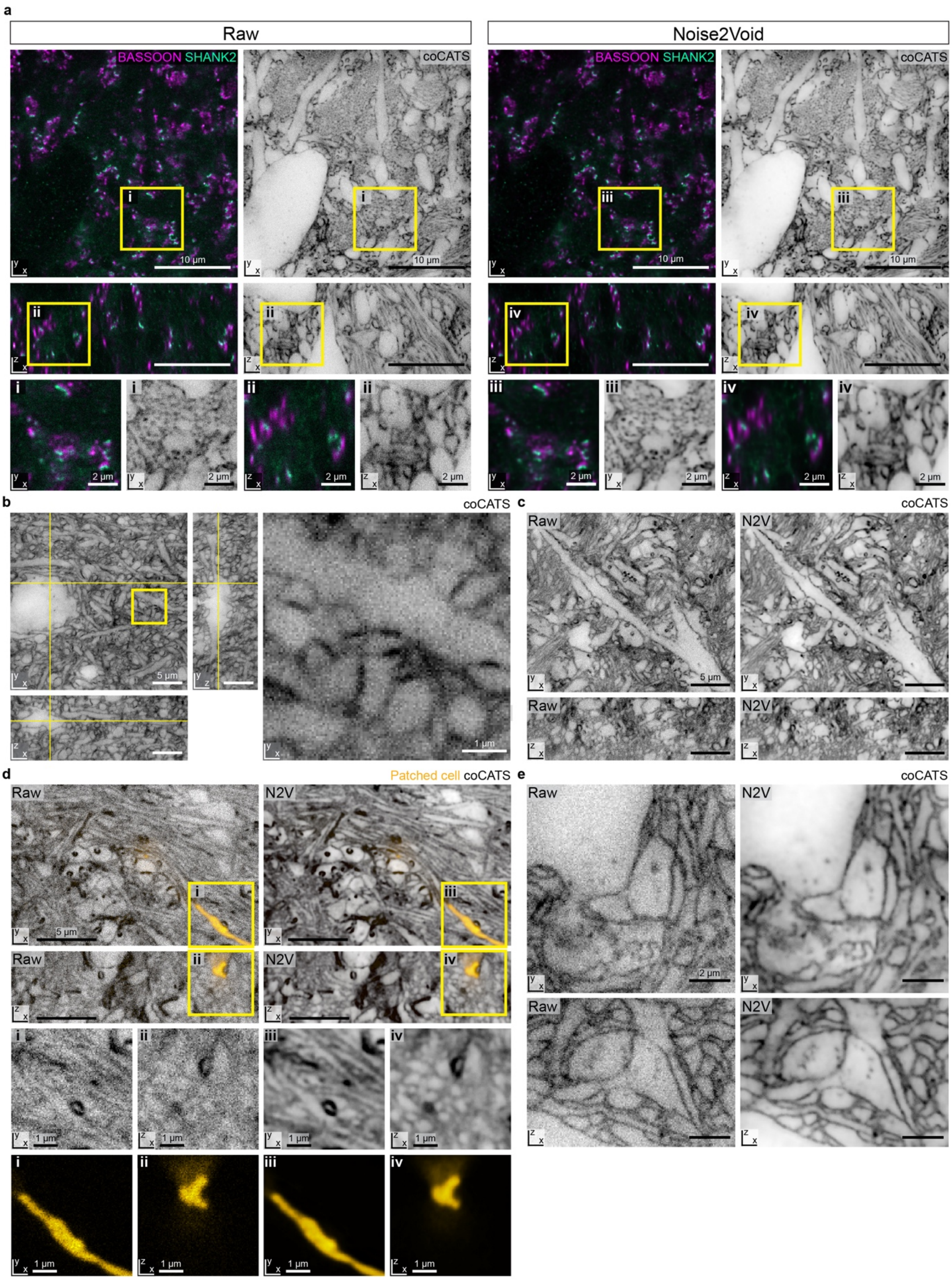


**Supplementary Fig. 7| Comparison of raw vs. Noise2Void (N2V)-denoised imaging data.**

**a,** Single *xy*- and *xz*-section, as well as zoomed views (i-iv) from one of the three volumetric imaging datasets used for MFB- and pSCR-segmentation in Fig. 2. Raw (*left*) and N2V-denoised (*right*) data of BASSOON (magenta, confocal), SHANK2 (turquoise, *z*-STED) and coCATS (grey, *z*-STED). **b,** CoCATS raw data (*z*-STED) corresponding to the dataset in Fig. 3a,b, showing the same *xy*-, *xz*-sections and zoomed view of the boxed region. **c,** Raw (*left*) and N2V-denoised (*right*) single *xy*- and *xz*-section of the volumetric coCATS imaging dataset (*z*-STED) displayed in Fig. 3g, used for reconstruction of the input field of a CA3 pyramidal neuron proximal dendrite. **d,** Single *xy*- and *xz*-sections with zoomed views of the boxed regions (i-iv), corresponding to one of the volumetric imaging datasets used for the characterization of a DG granule cell output field (Fig. 4 c,d), displaying coCATS (grey, *z*-STED) and intracellular label (yellow, *z*-STED) **e,** Single *xy*- and *xz*-sections of the volumetric imaging dataset of a human cerebral organoid (coCATS, *z*-STED) displayed in Fig. 6d without (*left*) and with N2V.

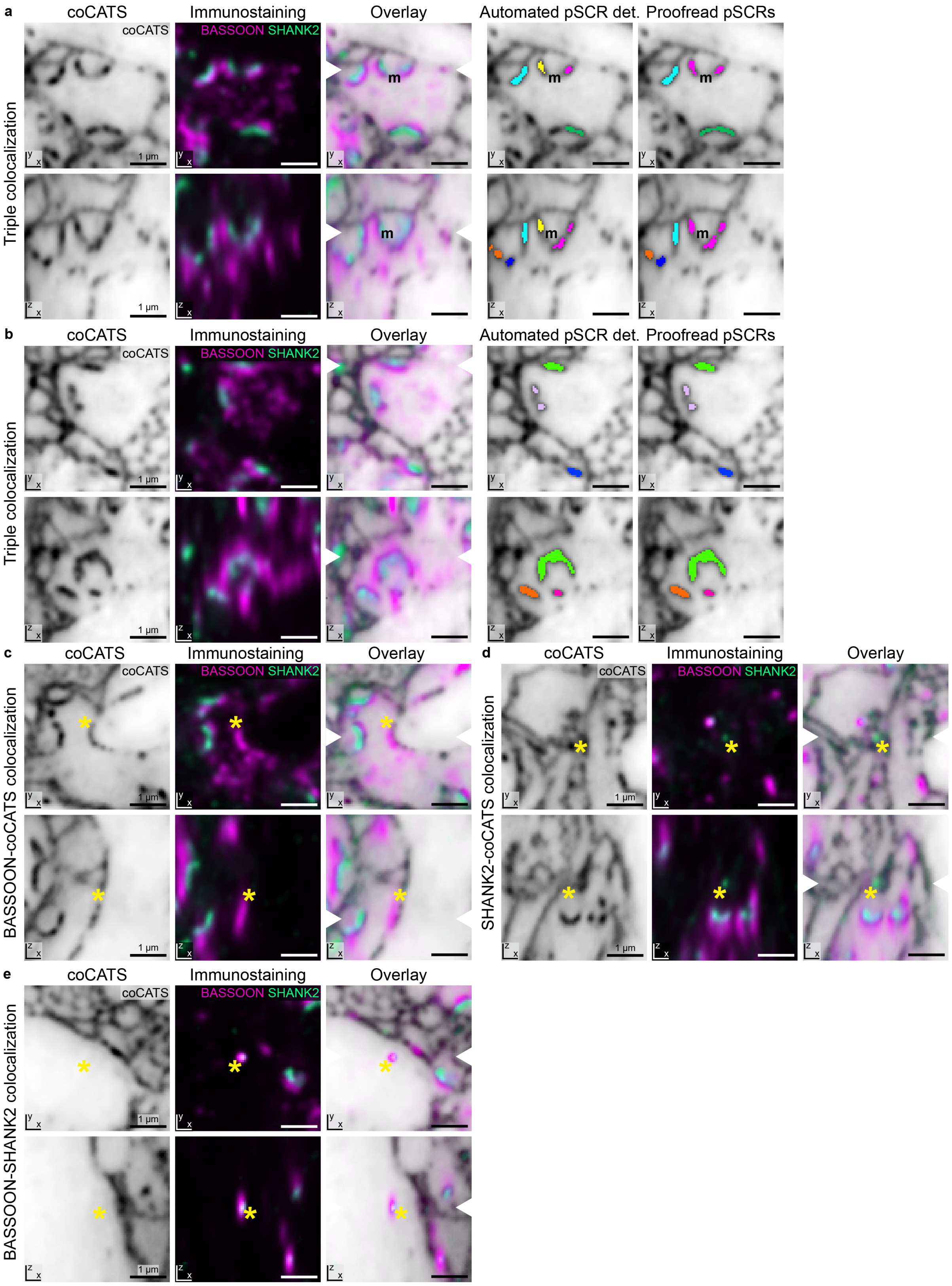


**Supplementary Fig. 8| Specificity of automated synapse detection.** Examples of colocalization types between BASSOON (magenta, confocal), SHANK2 (turquoise, *z*-STED), and high intensity coCATS features (grey, *z*-STED) taken from the 3 imaging volumes used for reconstruction and quantification of single-MFB properties in Fig. 2. Single *xy*- and *xz*-sections of coCATS, immunostaining, and overlay, with all channels denoised by N2V. White arrowheads in the overlays indicate positions of the orthogonal views. **a,b,** Orthogonal views for two examples with various instances of BASSOON, high intensity coCATS, and SHANK2 triple colocalization, resulting in pSCR detection (det.) by the automated pipeline. PSCR instance segmentations are displayed multi-color and shown as raw output of the automated segmentation and after manual proofreading of the 3D data. m: example of segments that were merged during proofreading. Scale bars: 1 µm. **c,** Example of colocalization (asterisk) of BASSOON and coCATS feature without SHANK2 staining present, likely corresponding to an inhibitory synapse. **d,** Example of colocalization (asterisk) of SHANK2 and coCATS feature, without BASSOON staining present. Note that SHANK2 signal is weak compared to local surrounding, hinting towards non-specific antibody binding close to a region of high intensity in the coCATS channel. **e,** Example of colocalization (asterisk) of BASSOON and SHANK2 in a region without high intensity coCATS feature. Here, BASSOON and SHANK2 are closely associated and the signal shapes approximate the microscope point spread functions for confocal and STED. This is thus likely a result of non-specific binding of secondary antibodies. In the *xz*-view, the difference in resolution between STED (green) and confocal (magenta) is particularly evident. Cases of double detection, as displayed in c-e, are not classified as synapses by our pipeline for automated pSCR segmentation.

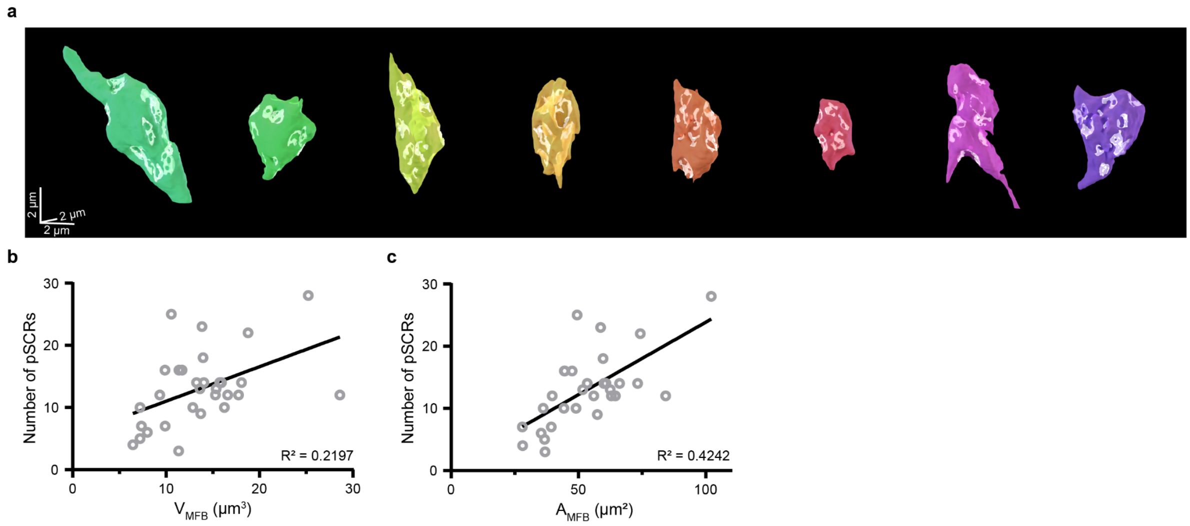


**Supplementary Fig. 9| Additional reconstructed MFBs and quantifications. a,** Eight additional reconstructed MFBs with pSCRs (white shaded areas) representing together with the 22 reconstructed boutons in Fig. 2b the total of 30 MFBs reconstructed and quantified in Fig. 2b-g. **b,c,** Number of pSCRs associated with individual boutons as a function of MFB volume ($V_{MFB}$) and MFB surface area ($A_{MFB}$), including linear regression ($V_{MFB}$: Pearson correlation coefficient r=0.4687, CI: 0.131-0.709, p=0.009; $A_{MFB}$, r=0.6513, CI: 0.380-0.819, $p<0.0001$). The correlation of MFB volume or MFB surface area with the number of associated pSCRs is lower than with the pSCR area per bouton ($A_{pSCR/MFB}$, Fig. 2f,g), indicating that the number of transmission sites may be less stringently controlled than the overall area of an MFB devoted to synaptic transmission. Note, however, that any merges of adjacent pSCRs due to finite resolution would be directly reflected in pSCR count but less so in pSCR covered surface area.

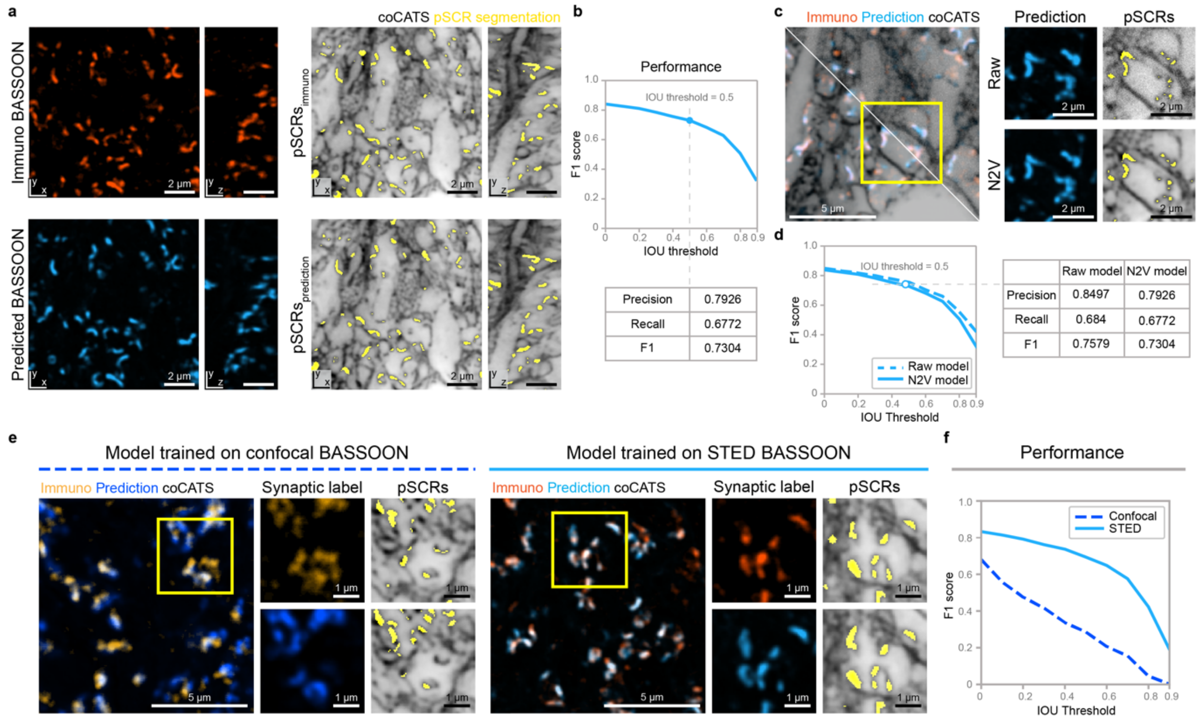


**Supplementary Fig. 10| Validation of deep-learning-based prediction of synaptic marker location and pSCR segmentation from coCATS structural data.** CATS datasets were acquired after *in vivo* microinjection and combined with immunostaining for BASSOON (both channels recorded volumetrically with a *z*-STED pattern for predominantly axial resolution increase, unless specifically noted). Data were denoised with Noise2void (N2V), unless otherwise noted. Validation datasets were not included in training of the convolutional neural network. **a,** (*Left*) Comparison of *xy*- and *xz*-planes for immunostained BASSOON (*top*) and deep-learning-predicted BASSOON location (*bottom*). (*Right*) CoCATS channel (grey) with corresponding pSCR segmentations (yellow) based on immunostained BASSOON (*top*) and predicted BASSOON (*bottom*). For this comparison, automated pSCR segmentations without proofreading were used both for immunostained and predicted BASSOON. **b,** F1 score (incorporating precision and recall) for comparing pSCR segmentation based on predicted BASSOON signal against pSCR segmentation based on immunostained BASSOON signal as a function of intersection over union threshold (IOU threshold). F1, precision, and recall in tabular form at IOU threshold=0.5, indicating overall high performance of the deep-learning-based model for pSCR prediction. **c,** Comparison of prediction of BASSOON signal and pSCR segmentation by two models trained on denoised vs. raw coCATS data. (*Left*) Immunostained (orange) and predicted (blue) BASSOON with either raw or denoised CATS data. Magnified views of predicted BASSOON and pSCR segmentations for raw (*top*) and N2V (*bottom*) CATS data. Segmentation results are similar. **d,** Similar performance is also evident in the F1 score as a function of IOU threshold and in the table of performance parameters, benchmarking the prediction of synaptic marker location by models trained on raw CATS data and on N2V CATS data against the immunostaining. **e,** Comparison of network models for prediction of BASSOON signal trained on super-resolved CATS (*z*-STED) plus super-resolved BASSOON immunostaining (*z*-STED) vs. training on super-resolved CATS (*z*-STED) plus confocal immunostaining data. (*Left*) Training based on confocal BASSOON with magnified view of the boxed region for immunostained BASSOON, predicted BASSOON signal, and corresponding pSCR segmentations. (*Right*) Training using super-resolved BASSOON. **f,** F1 score as a function of IOU threshold for confocal vs. STED models. The performance is markedly lower when using confocal BASSOON data, showing that the higher resolution in STED imaging is necessary to train a model to faithfully predict synaptic marker location and localize pSCRs in the deep-learning-based segmentation pipeline.

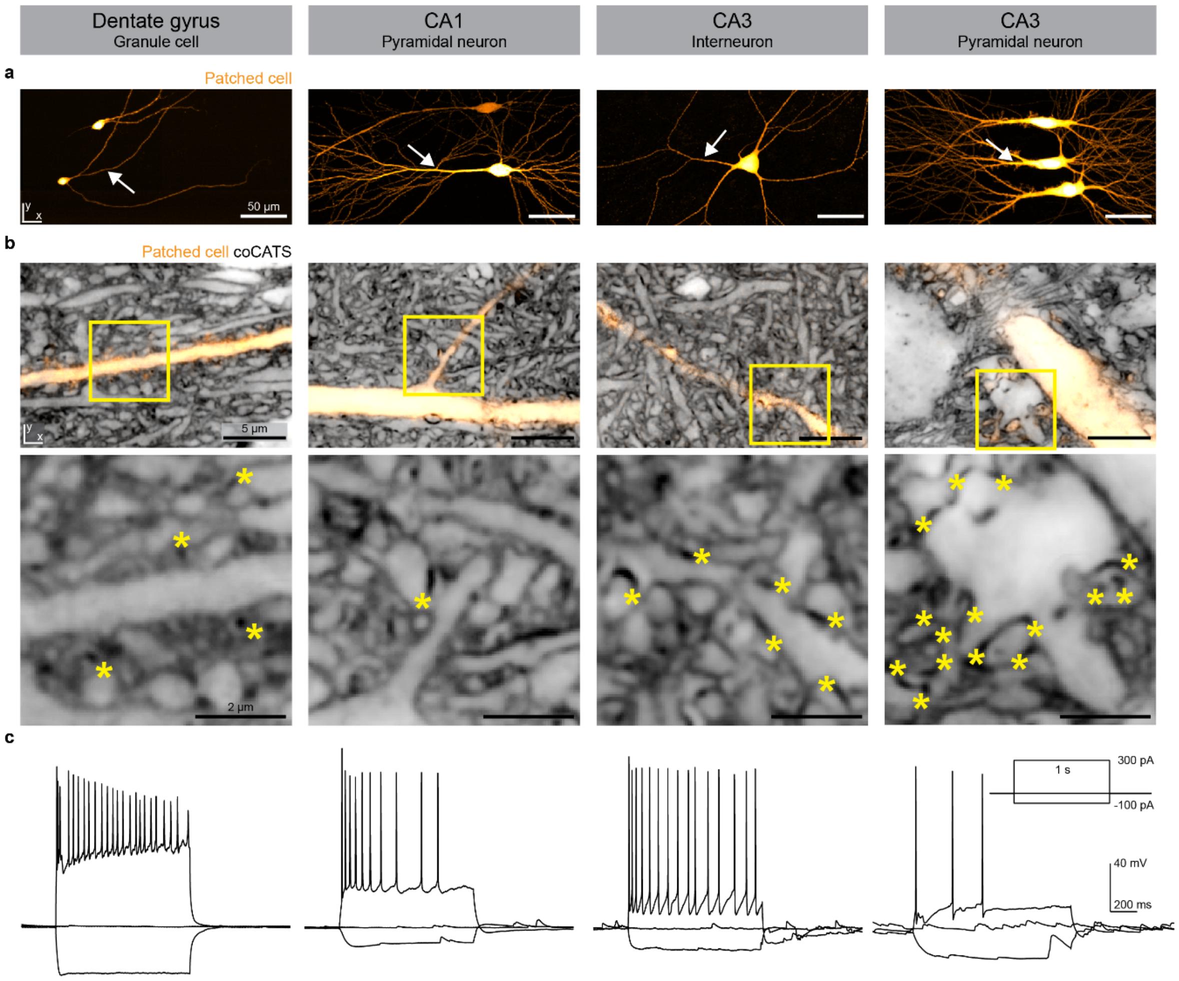


**Supplementary Fig. 11| Combined structural and functional characterization with coCATS and single-cell patch-clamp experiments in various neuronal cell types.** Neurons in organotypic hippocampal slice cultures were whole-cell patch-clamped, recorded and dye-filled with Lucifer yellow before coCATS labeling, immersion-fixation and imaging. **a,** Maximum intensity projection (MIP) overview images of various patched cells (DG granule cell, CA1 pyramidal neuron, CA3 interneuron, CA3 pyramidal neurons) acquired with a confocal microscope. Scale bars: 50 µm. **b,** (*Top*) Single *xy*-planes of volumetric imaging datasets showing close-ups (white arrows in panel a) of the same cells (yellow, confocal, N2V) in their tissue context revealed by coCATS (grey, *z*-STED, N2V). (*Bottom*) Zoomed views of boxed regions. CoCATS visualizes the local environment of the cells, as well as the synaptic connectivity profile via pSCRs (asterisks). This information is useful for characterizing cell types and their connectivity also in the absence of an immunostaining. Scale bars: 5 µm. **c,** Unique action potential phenotypes upon current injection into the patched cells provide complementary information for cell type identification and single cell characterization.

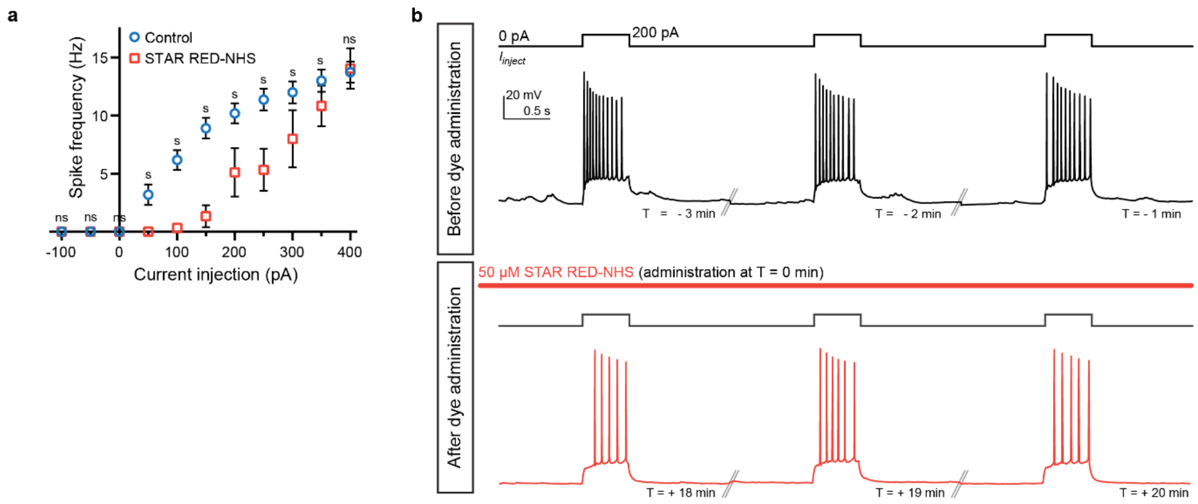


**Supplementary Fig. 12| Neuronal activity during and after coCATS incubation. a,** Action potential frequency as a function of current injection in CA1 pyramidal neurons in 15-16 DIV organotypic hippocampal brain slices without coCATS labeling (Control) and after 25 min incubation with 50 µM STAR RED-NHS. While viability and maximum firing rate were not affected (ns=non-significant, p-value>0.05, unpaired t-test), lower spiking frequency is observed at lower injection currents in coCATS-incubated samples (50-350 pA, s=significant, p-value<0.05, unpaired t-test). This effect likely results from an increase in network activity during dye incubation. Such increased network activity can commonly be observed in hyperconnected organotypic cultures. **b,** Firing profile of an example CA3 pyramidal neuron in response to current injection in an organotypic hippocampal brain slice at 3 time points spaced by 1 minute before (*top*) and at 3 time points after application of the CATS label (*bottom*), spaced by 1 minute and starting 18 minutes after label application. CATS labeling does not prevent cell firing which would be indicative of cell damage.

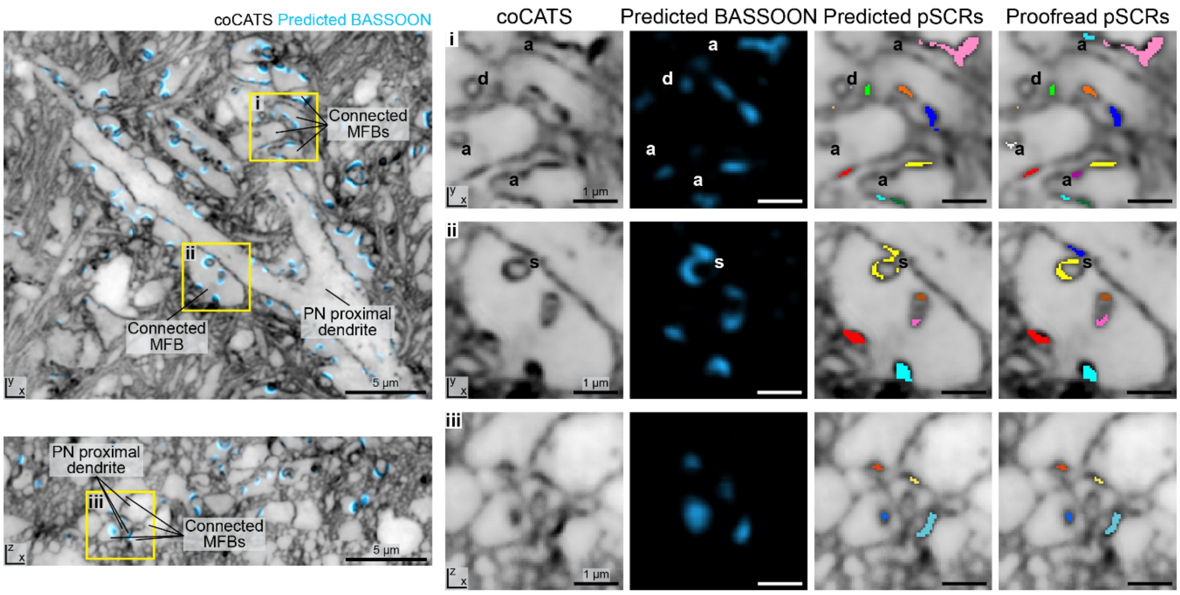


**Supplementary Fig. 13| Deep-learning-based identification of synapse location in coCATS labeled organotypic hippocampal slice cultures.** CA3 pyramidal neuron (PN) proximal dendrite as shown in Fig. 3g-i with synaptically connected mossy fiber boutons (MFB) identified via the presence of pSCRs. (*Left*) Single *xy*- and *xz*-planes of near istropically super-resolved coCATS (N2V) and BASSOON signal predicted by the deep learning algorithm (blue). (*Right*) Zoomed views of the boxed regions showing the CATS and BASSOON prediction channels separately, as well as automatically predicted and manually proofread pSCR instance segmentations (both multi-colored). Specific examples of proofreading operations include adding (a), deleting (d), splitting (s), and merging (not shown) segments as indicated.

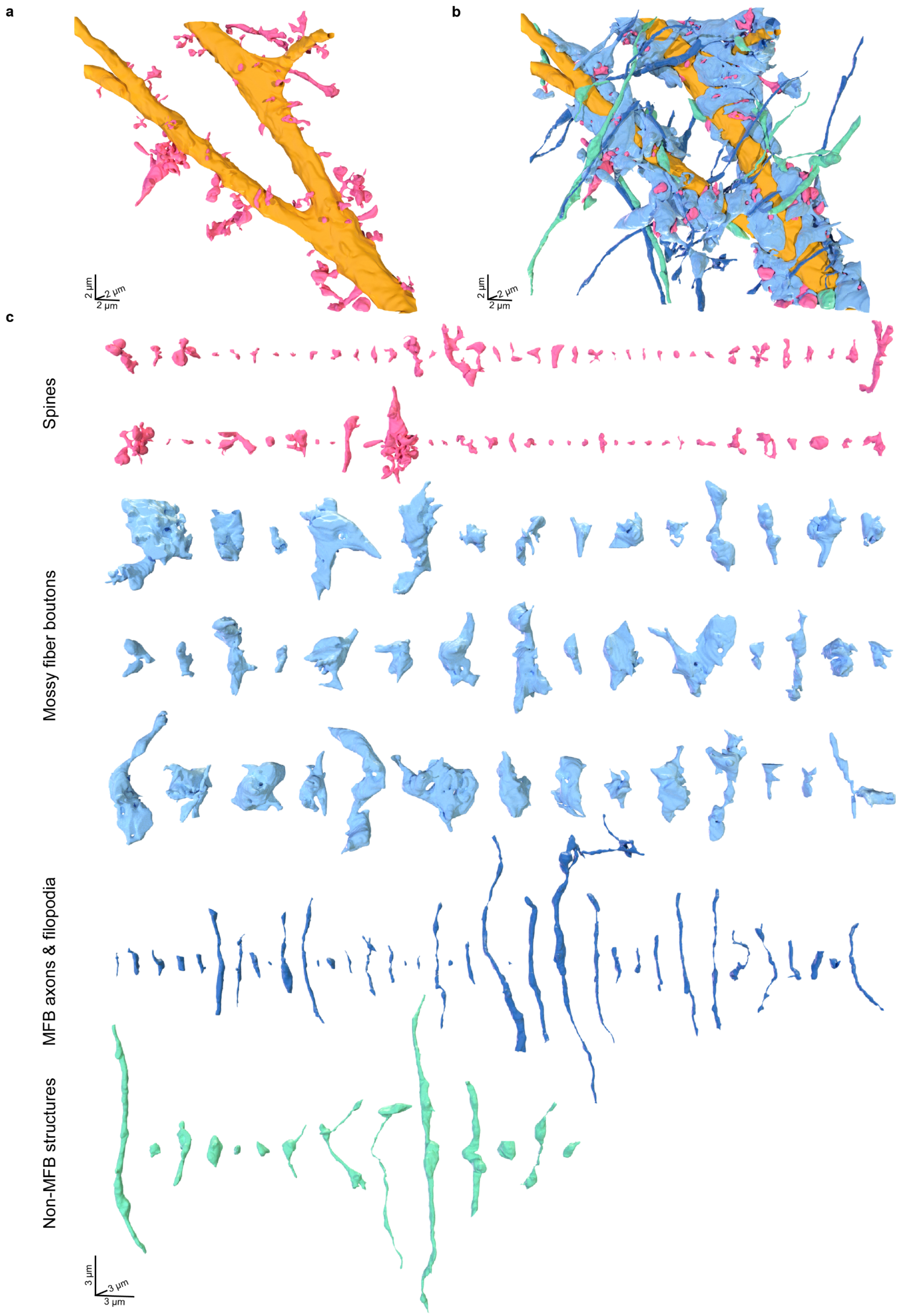


**Supplementary Fig. 14| Structural characterization of the local input field in a CA3 pyramidal neuron proximal dendrite. a,** 3D-rendering of the CA3 pyramidal neuron proximal dendrite in Fig. 3g based on coCATS data. The dendritic shaft is colored in gold, spines are labeled in magenta. **b,** 3D-rendering of the same dendrite as in a, with associated cellular structures color-coded by identity, as inferred from morphology: MFBs (light blue), axons and filopodia of MFBs (dark blue), structures in synaptic contact with the main dendrite, not identifiable as MFB-related structures (turquoise). **c,** 3D-renderings of all structures reconstructed in b.

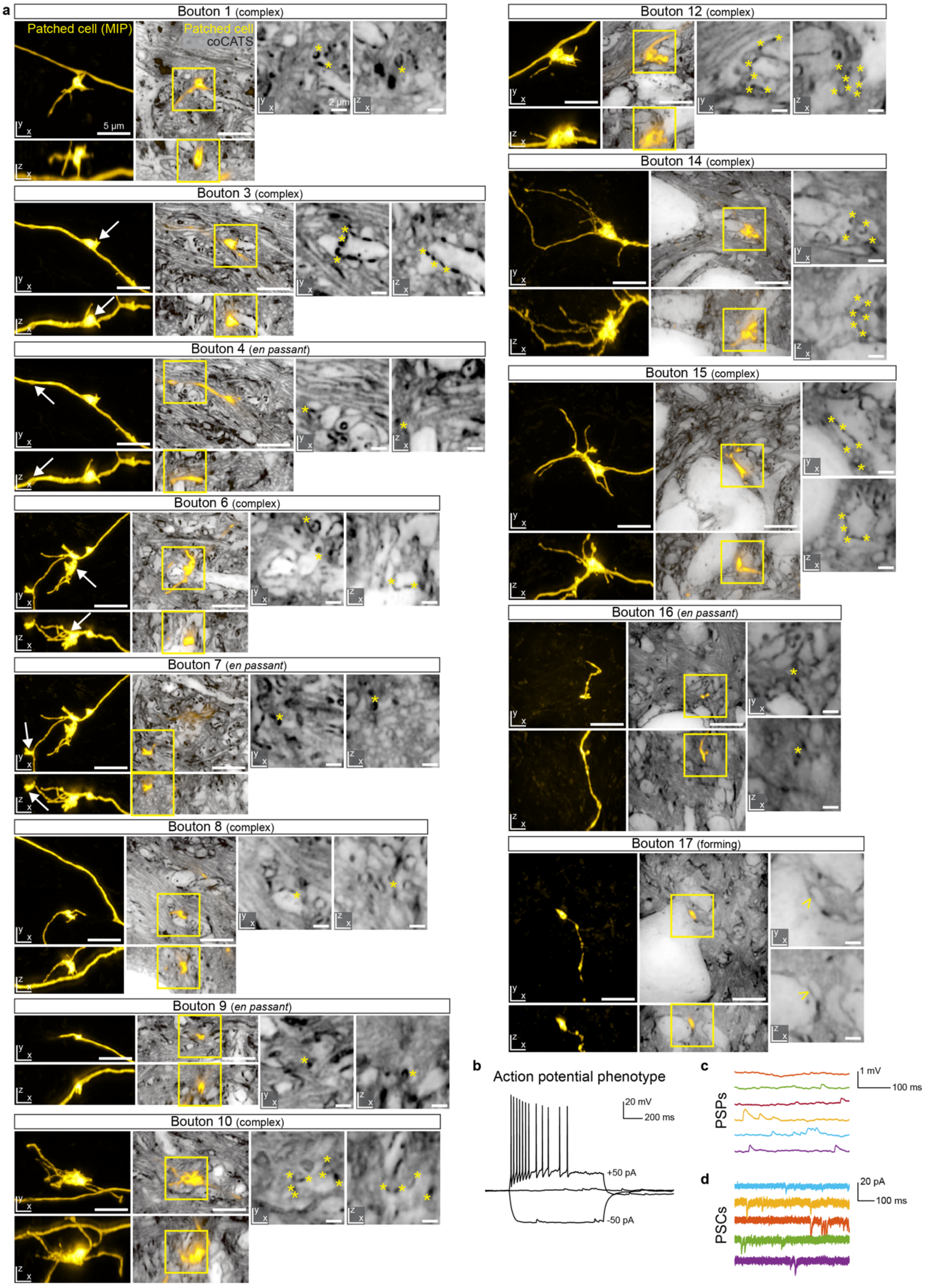


**Supplementary Fig. 15| Visualization of DG granule cell output field. a,** Tissue architecture around MFBs for all synaptic outputs marked in Fig. 4a in addition to the ones displayed in Fig. 4c. (*Left*) Maximum intensity projections (MIP) of positively labeled neuron (yellow, *z*-STED, N2V). (*Middle*) Single *xy*- and *xz*-planes from volumetric measurement of surrounding tissue architecture by coCATS (*z*-STED, grey, N2V) with overlaid positive label of the single neuron. (*Right*) Zoomed views as indicated by the boxes showing the CATS channel alone with asterisks indicating pSCRs. White arrows in the left panels indicate the synaptic bouton that is displayed in the CATS panels for cases with more than one bouton in the field of view. At bouton 17, small yellow arrows mark the end-point of the axon without a clear pSCR present (potentially developing synapse). Scale bars, overview images: 5 µm; zoom-ins: 2 µm. **b,** Action potential phenotype of the same DG granule cell, elicited by current injection during whole cell patch clamp recording. **c,** Spontaneous post-synaptic potentials (PSPs) and post-synaptic currents (PSCs) in the same neuron.

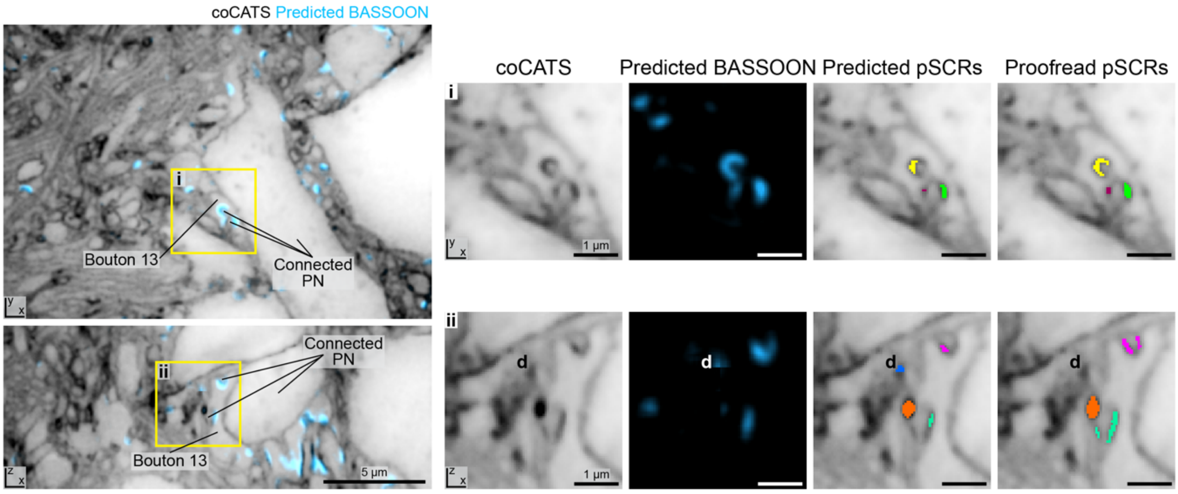


**Supplementary Fig. 16| Deep-learning-based identification of synapse location in coCATS labeled organotypic hippocampal slice cultures.** Imaging volume used for reconstructing the local output of bouton 13 of the DG granule cell in Fig. 4e. The location of bouton 13 of the positively labeled DG granule cell and a pyramidal neuron constituting its main post-synaptic partner (labeled 'Connected PN') are indicated. Scale bars, overviews: 5 µm; zoomed views: 1 µm. (*Left*) Single *xy*- and *xz*-planes of near-istropically super-resolved coCATS (N2V) and BASSOON signal predicted by our deep learning algorithm (blue). (*Right*) Zoomed views of the boxed regions showing the CATS and prediction channels separately, as well as automatically predicted and manually proofread pSCR instance segmentations (both multi-colored). In this specific example, a deletion operation (d) was performed during proofreading.

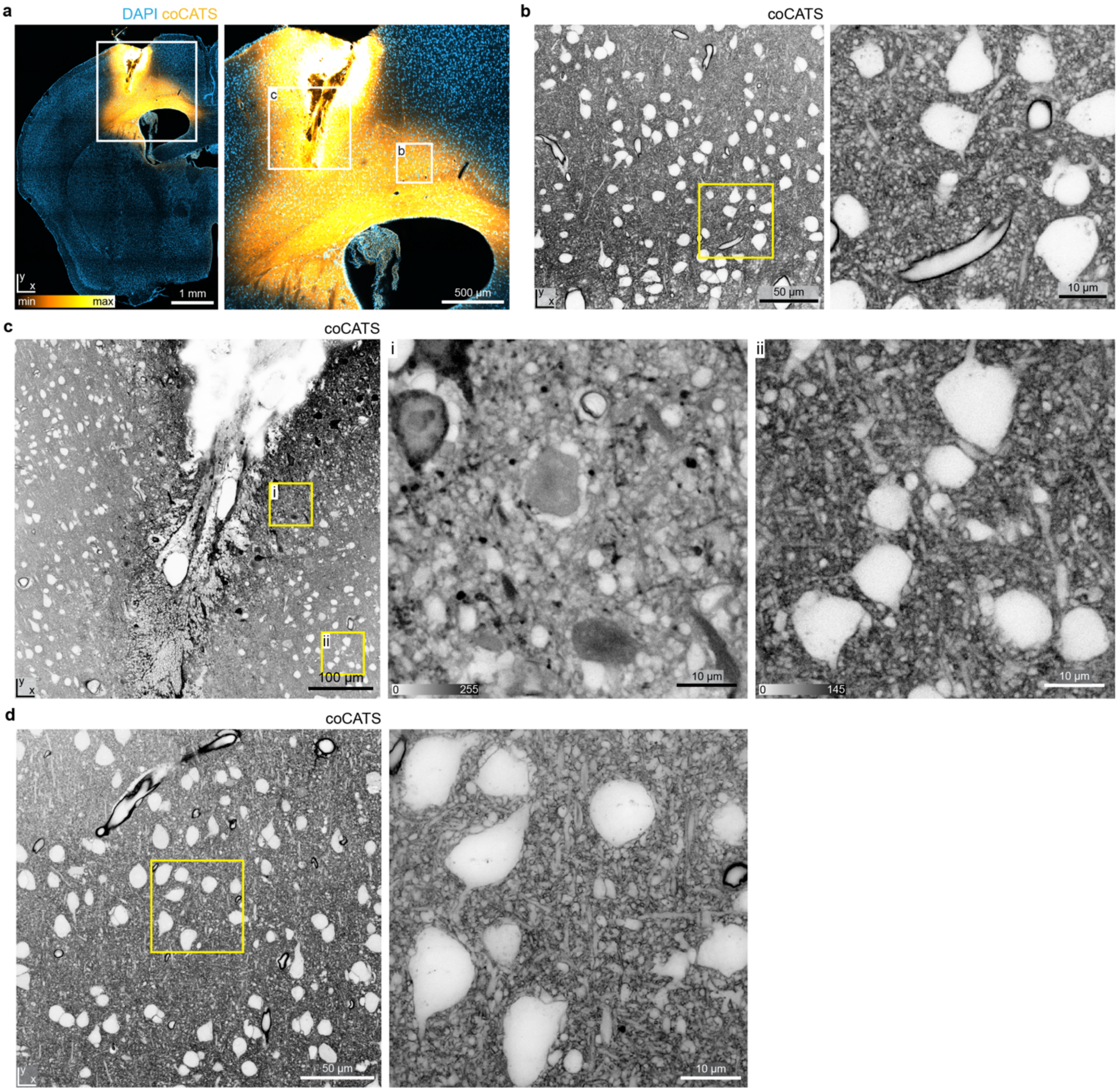


**Supplementary Fig. 17| Targeted delivery of coCATS label to a specific brain region by *in vivo* microinjection.** Brain areas of interest can be labeled by *in vivo* microinjection into the tissue close to the region of interest. **a,** Tile scan of a coronal brain section after *in vivo* microinjection into the somatosensory cortex. The injection site, which displays local damage, is clearly visible. Higher resolution image of the area indicated by the white rectangle shows that the damage is spatially limited, such that cortical structures in proximity to the injection site can be studied. CoCATS intensity lookup table (yellow) is not inverted. Data was acquired on a spinning disc confocal microscope. **b,** Enlarged view of a cortical area spatially separated from the injection site. High labeling intensity, but no tissue damage, is visible in this dataset. Injection of coCATS label at ~0.2-1.0 mm from the region of interest yields high contrast labeling. The right panel is a magnified view of the boxed region. **c,** Enlarged view of a region comprising the injection site with two zoomed views, one in the damage region in immediate proximity to the injection site and one in the nearby region of intact tissue. Local damage is visible by the presence of erythrocytes and big voids in the overview image. (i) Enlarged view of a region in immediate vicinity of the injection site. Cellular structures appear swollen and disorganized. Damaged cells are highly labeled (black), because the dye binds strongly to the protein-rich intracellular environment. Tissue structure cannot be appreciated as many structures have taken up labeling compound. (ii) Enlarged view of a region ~250 µm away from the injection site. No strongly labeled cellular structures are visible, and tissue organization, including cell bodies and processes, is visible. Images were acquired with a confocal microscope. **d**, Another region in the same coronal section. Overview confocal image (*left*) and zoomed view of the yellow-boxed region (*right*) acquired with STED microscopy show well-preserved tissue structure. The STED image was recorded with a power distribution between *xy*- and *z*-STED patterns of 85/15. Raw data.

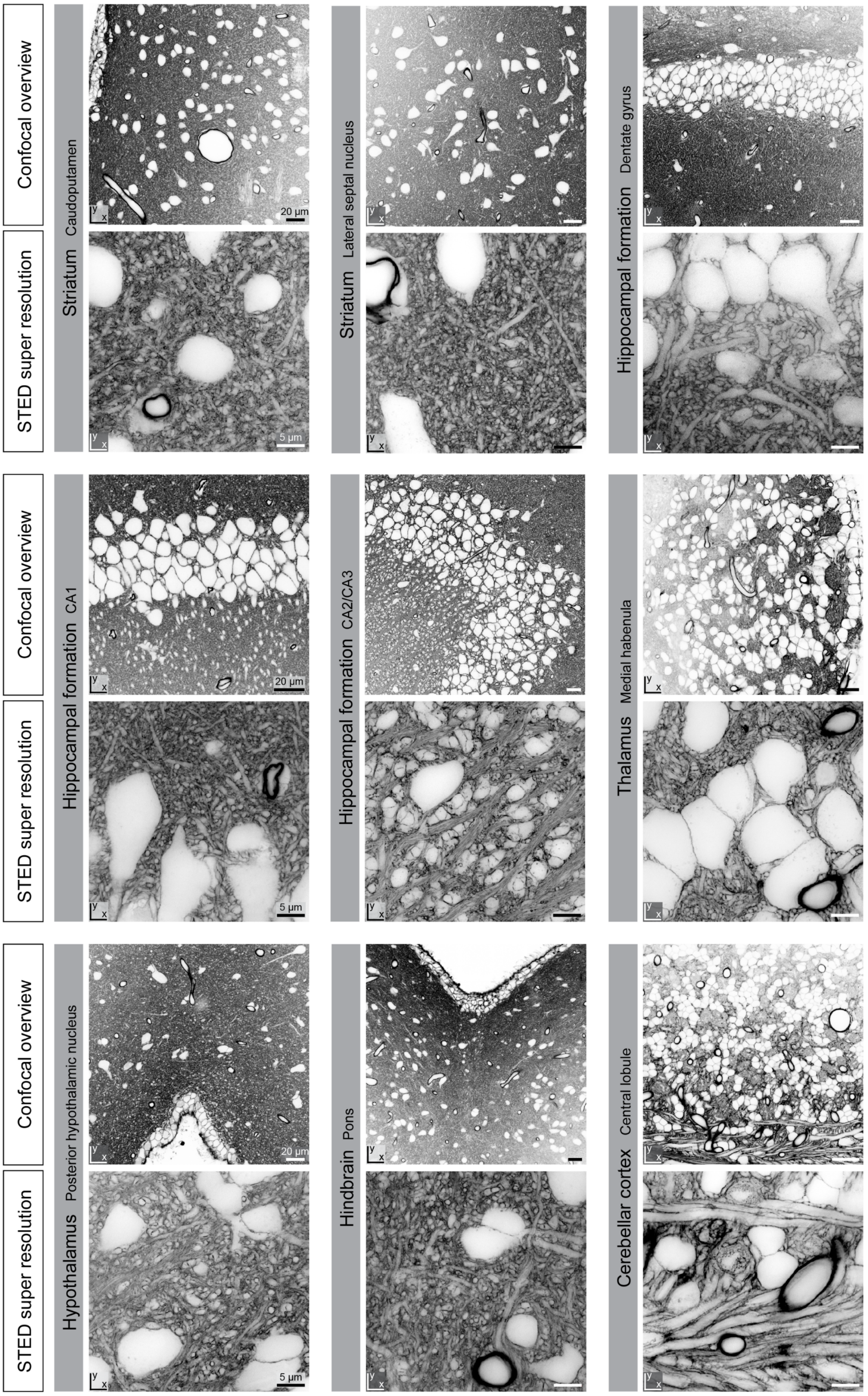


**Supplementary Fig. 18| Tissue organization in diverse brain regions.** CoCATS in various brain regions after *in vivo* microinjection into the lateral ventricle or cortex. (*Top*) Confocal overview images. Scale bars: 20 µm. (*Bottom*) Higher magnification STED images with lateral resolution increase (*xy*-STED) from the same regions. Raw data. Scale bars: 5 µm

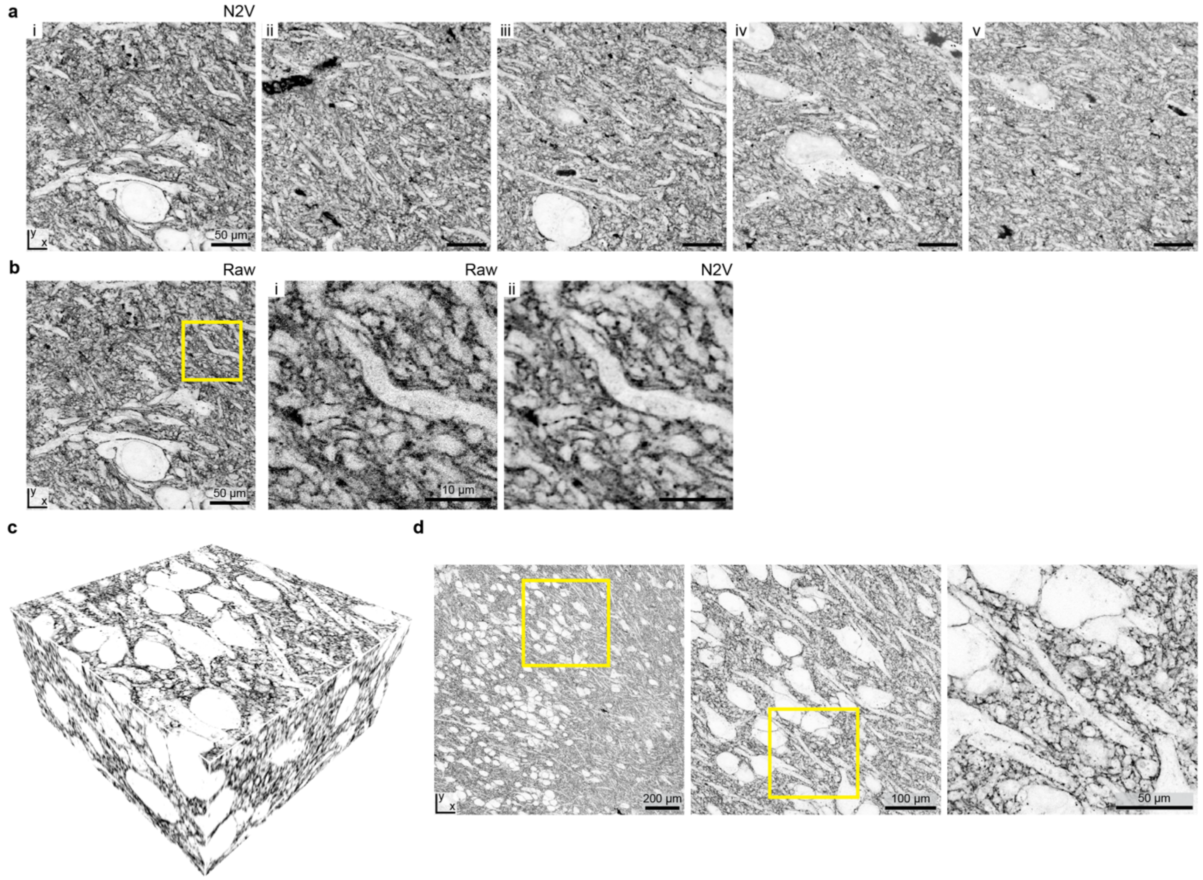


**Supplementary Fig. 19| CoCATS with ExM in organotypic hippocampal slice cultures. a,** En face views of the 5 example planes in Fig. 5b (denoised with N2V). Scale bars refer to tissue size after expansion. **b,** Raw data for the first slice, with zoom in the boxed region for (i) raw data and (ii) data after denoising with N2V. **c,** 3D-view of a 290 x 290 x 137 $\mu m^3$ imaging volume after ~4-fold expansion of an organotypic hippocampal slice culture with protein retention ExM. The tissue was coCATS labeled with NHS-$PEG_{12}$-biotin, immersion-fixed, hydrogel-embedded, mechanically homogenized by proteolytic digestion, and expanded. Post-expansion readout was performed with fluorophore-labeled streptavidin. **d,** Overview image of a single *xy*-plane of the same sample and progressive zoom-ins as indicated by the rectangles. Data was acquired with a confocal microscope. Scale bars refer to sample size after expansion.

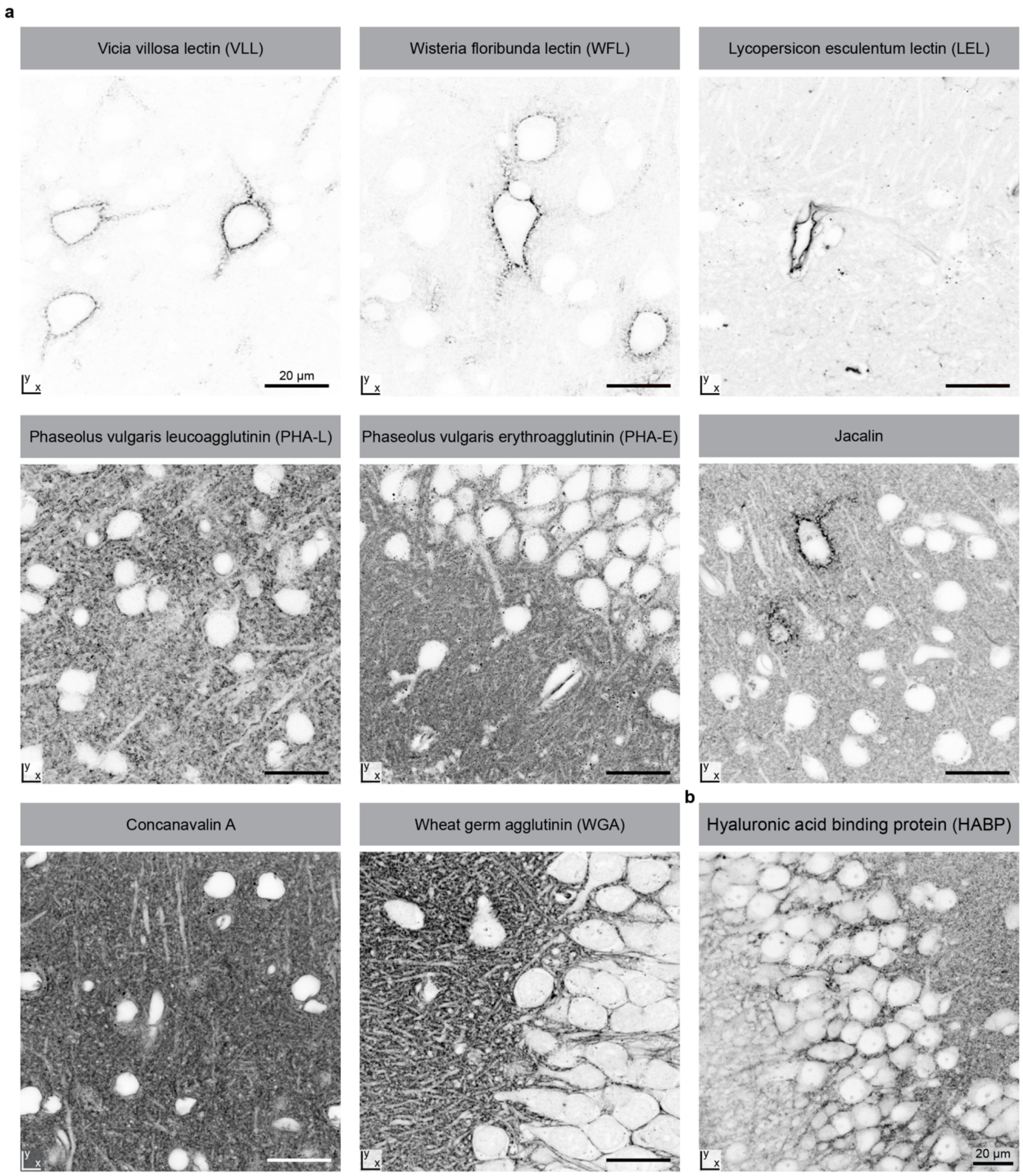


**Supplementary Fig. 20| Screening for affinity binders to reveal tissue architecture with resident CATS (rCATS) in rodent brain. a,** Binding pattern of various lectins in previously fixed adult mouse brain. (*Top, left*) *Vicia villosa* lectin (VVL) and (*top, center*) *Wisteria floribunda* lectin (WFL) binding patterns are restricted to perineuronal nets associated with a sparse subset of cells (cortex). These are thought to bind to terminal N-acetylgalactosamine linked to serine/threonine and galactose, respectively. (*Top, right*) Lycopersicon esculentum lectin (LEL) faithfully depicts blood vessels by binding to specific N-glycans (hippocampus). (*Middle, left*) *Phaseolus vulgaris* leucoagglutinin (PHA-L) and (*middle, center*) *Phaseolus vulgaris* erythroagglutinin are two members of the same lectin family that resulted in a grainy extracellular staining pattern in the adult mouse cortex (PHA-L) and dentate gyrus (PHA-E). (*Middle, right*) Jacalin binds to O-glycosidically linked oligosaccharides, leading to a strong labeling of perineuronal nets, as well as a weak depiction of the extracellular environment. (*Bottom, left*) Concanavalin A recognizes α-mannose on oligosaccharides. This results in a homogeneous labeling of the extracellular space, here shown in cortex, but also of intracellular structures, including the nuclear envelope. (*Bottom, middle*) Wheat germ agglutinin (WGA) delineates cell bodies and cellular processes by binding to N-acetylglucosamine and sialic acid abundant in the extracellular matrix and on cell surfaces, as shown here for hippocampus. Scale bars: 20 µm. **b,** Hyaluronic acid was labeled via fluorescently labeled hyaluronic acid binding protein (HABP) in the mouse hippocampus. Cellular outlines are visible, but the labeling is not homogeneous. All images were acquired with a confocal microscope. Intensity lookup tables are inverted.

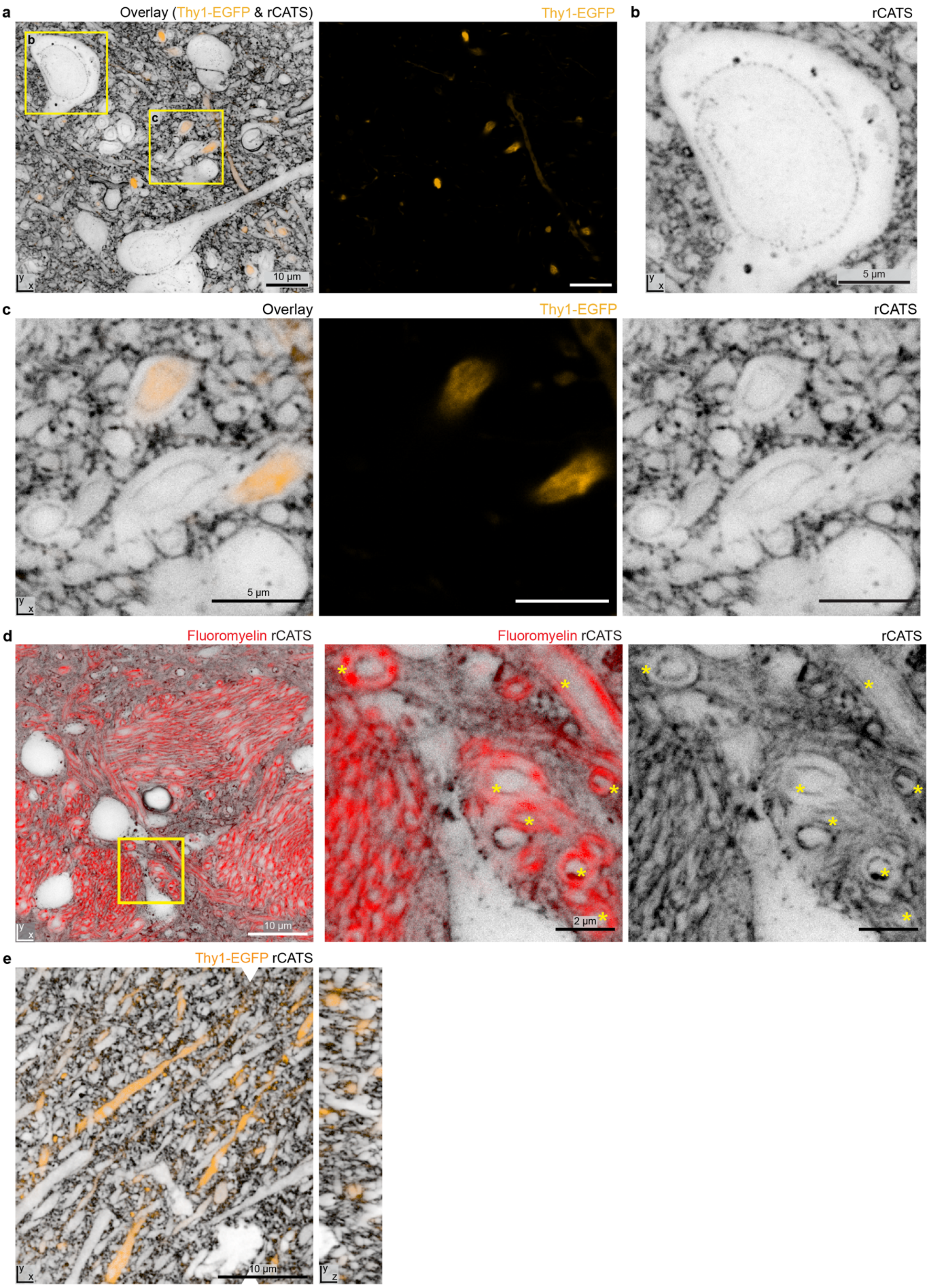


**Supplementary Fig. 21| Identification of myelination and nuclear pores by rCATS. a,** rCATS labeling in the hypothalamus of a perfusion-fixed adult Thy1-EGFP+ mouse. A sparse subset of neurons are labeled via cytosolic EGFP expression (orange, confocal), shown in the overlay with rCATS (grey, STED with power distribution of *z*-STED/*xy*-STED patterns of 80/20) and as a separate panel. **b,**

Magnified view of boxed region in a: Neuronal soma with nuclear envelope discernible in the rCATS channel due to binding of WGA to nuclear pore glycoproteins. **c,** Magnified view of second boxed region in a. Zoom-in on myelinated axons. Cytoplasmic expression of EGFP in a subset of neurons exclusively labels the axon but not the myelin sheath. In contrast, rCATS delineates the inner and outer borders of the myelin sheath. **d,** Fluoromyelin staining (red, confocal), as well as rCATS (grey, *xy*-STED), reveal myelinated axons in the hypothalamus of a perfusion-fixed adult mouse. Magnified view: myelinated structures delineated by rCATS co-localize with Fluoromyelin (examples indicated by asterisks). **e,** STED at near-isotropic resolution of rCATS labeling (grey, *z*-STED) in the cortex of a perfusion-fixed adult Thy1-EGFP+ mouse (EGFP: orange, confocal). White arrowheads indicate position of *yz*-view.

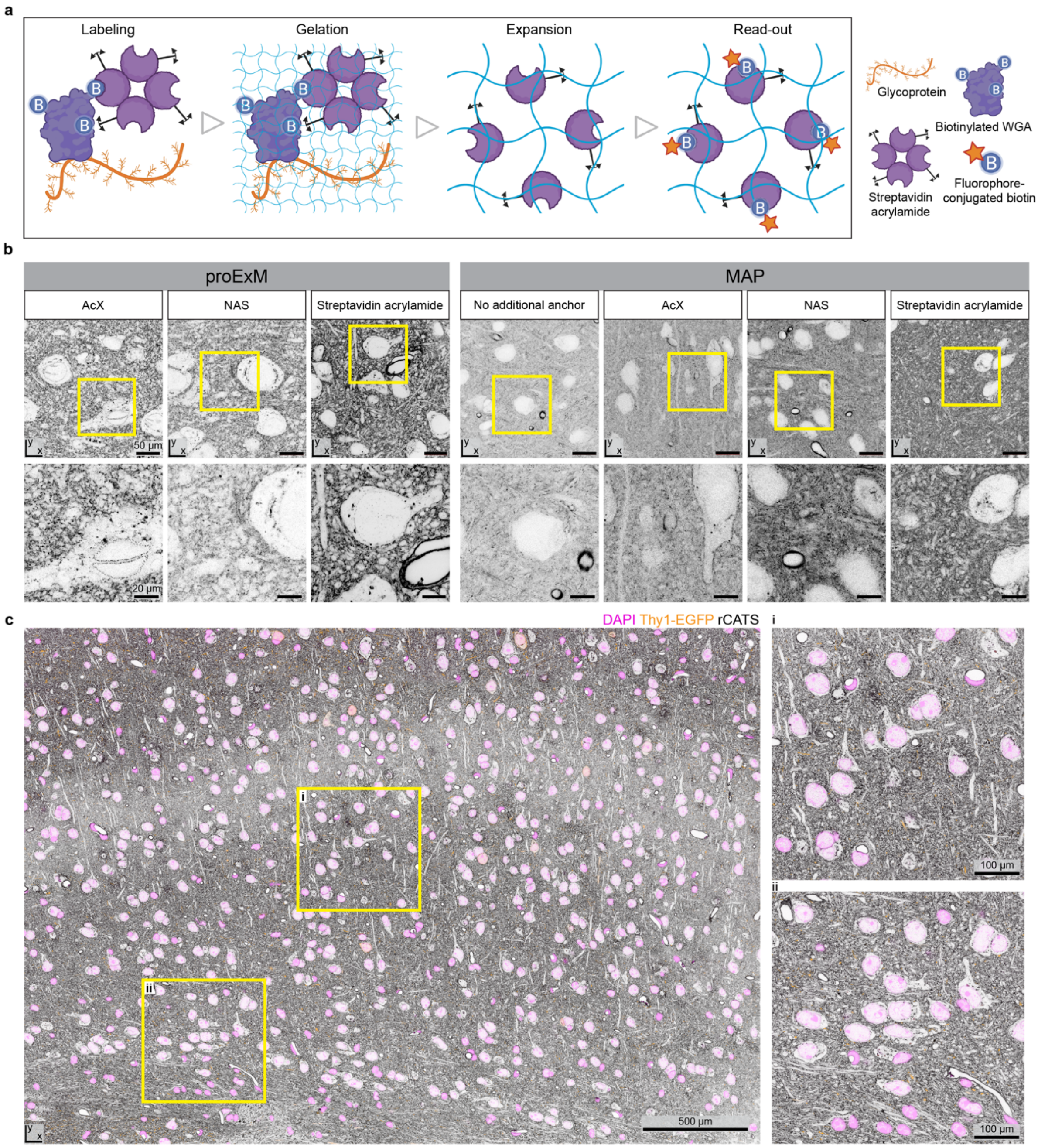


**Supplementary Fig. 22| Pipeline for signal retention with rCATS in expansion microscopy. a,** Schematic of rCATS expansion pipeline ensuring retention of WGA signal. Biotinylated WGA labels predominantly extracellularly located glycoproteins. To ensure that the WGA signal is retained in the hydrogel, the biotin on the WGA is targeted with streptavidin acrylamide. Upon gelation, streptavidin copolymerizes with the hydrogel via the acrylamide moiety. During the expansion procedure, polysaccharides and biotinylated WGA may get lost, but the anchored streptavidin remains in place and is read out with fluorophore-conjugated biotin post-expansion. Streptavidin labeling does retain its capacity to bind biotin after common homogenization procedures for disrupting tissue cohesiveness in ExM, including heat/chemical denaturation and enzymatic digestion. **b,** Retention of biotinylated WGA in slices from PFA-perfused mouse brain was tested with two common ExM strategies, protein-retention ExM (proExM) and magnified analysis of proteomes (MAP), using various retention strategies. Anchoring for proExM with acrylic acid N-hydroxysuccinimide ester (NAS) or Acryloyl-X (AcX) led to specific, but low intensity signal, likely due to the limited number of lysine residues on WGA that can be targeted by such an approach. In contrast, a highly specific and strong WGA labeling pattern was obtained when retaining WGA via streptavidin acrylamide and reading it out post-expansion with fluorophore-conjugated biotin. With the standard PFA/acrylamide based retention strategy in the MAP approach (no additional anchoring), the WGA signal was grainy and diffuse. Few structures, mainly blood vessels, were labeled strongly. Additional anchoring with AcX or NAS in MAP somewhat improved WGA-retention, but still resulted in an overall diffuse labeling pattern with strong labeling of blood vessels and putative myelinated processes. Diffuse labeling was also visible in cell bodies. Handing over WGA signal to streptavidin acrylamide improved the labeling quality further in MAP, but still resulted in inhomogeneities and aggregates, such that we opted for the proExM approach in this specific case. **c,** A slab of an adult Thy1-EGFP mouse cortex processed with the rCATS expansion pipeline, 4-fold expanded with proExM. (i-ii) Zoom-ins of the yellow boxed regions. Scale bars refer to sample size after expansion throughout. All images were acquired with a confocal microscope.

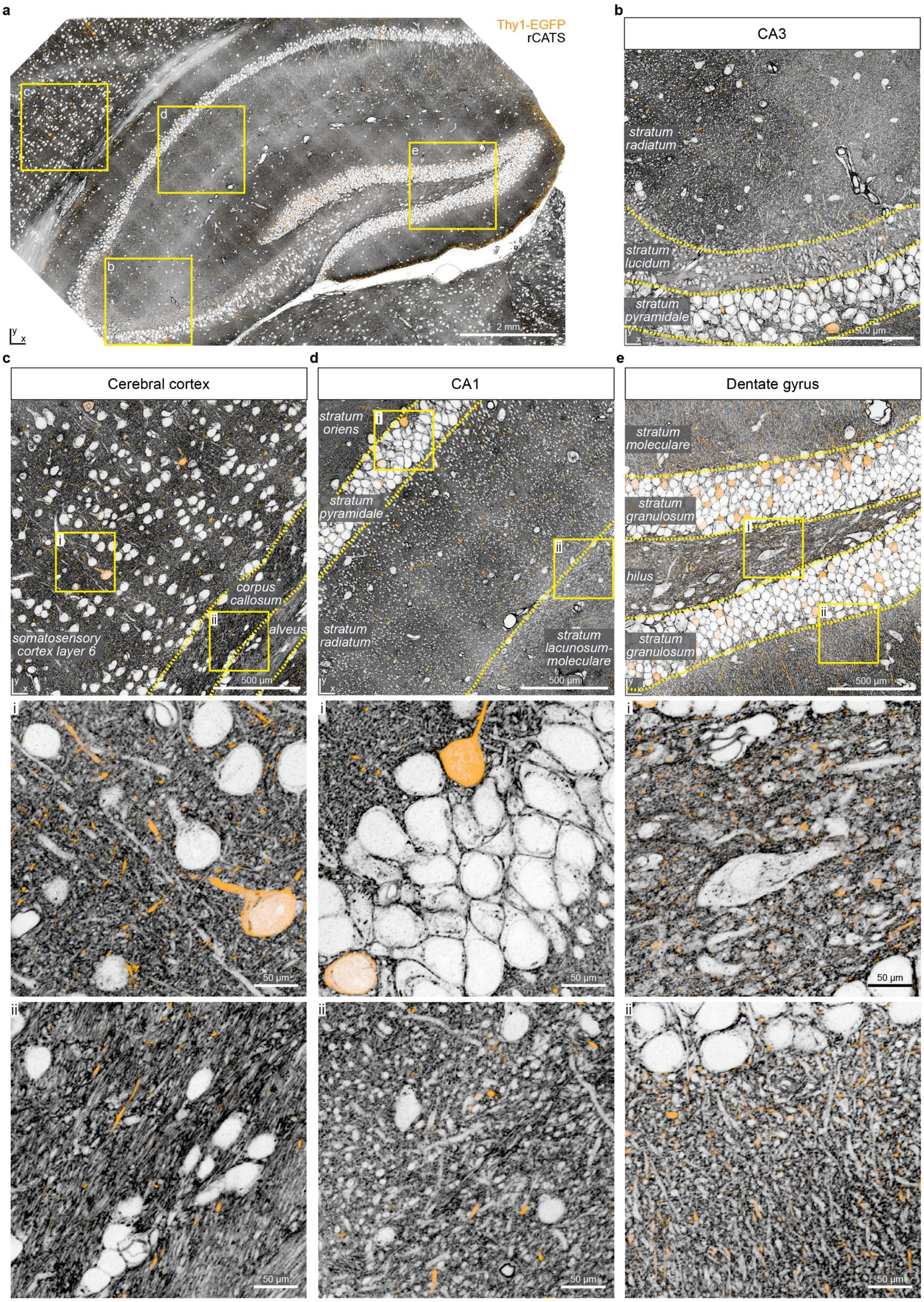


**Supplementary Fig. 23| Nervous tissue organization across scales revealed by rCATS and expansion microscopy.** Hippocampus and overlying cortex labeled with rCATS and expanded ~4-fold with protein retention ExM in a brain of an adult, perfusion-fixed Thy1-EGFP+ mouse, imaged with a spinning disc confocal microscope. A sparse subset of neurons is highlighted by cytosolic EGFP expression. **a,** Overview image of a coronal section of hippocampus and adjacent cortex. Boxes indicate the regions enlarged in the following panels. Scale bars refer to sample size after expansion throughout. **b**, Enlarged view of the CA3 region (same as in Fig. 5c). Hippocampal layers, including the CA3 *stratum radiatum*, CA3 *stratum lucidum*, and the CA3 pyramidal layer (*stratum pyramidale*) are clearly identifiable from the rCATS labeling. **c,** Enlarged view of the cerebral cortex with two further zoom-ins as indicated by the boxes. The somatosensory cortex layer 6 can be distinguished from the corpus callosum and the alveus by the differential organization of the tissue. The cortex contains many cell bodies and processes running perpendicular to the cortical surface, while the corpus callosum mainly consists of fiber tracts running parallel to the cortical surface. Sparse EGFP-positive cell bodies and neuronal processes can be found. **d,** Enlarged view of the CA1 area and two further zoom-ins as indicated by the boxes. The CA1 *stratum oriens*, CA1 *stratum pyramidale*, CA1 *stratum radiatum,* and CA1 *stratum lacunosum-moleculare* can be identified. **e,** Enlarged view in the dentate gyrus and two further zoom-ins. The organization of the DG blades, including the DG granule cell layer (*stratum granulosum*) surrounding the DG polymorph layer (hilus), as well as the DG molecular layer (*stratum moleculare*), can be appreciated. In the polymorph and molecular layers, sparse EGFP-positive structures, predominantly corresponding to DG axons and boutons, are visible.

### Supplementary Video captions

**Supplementary Video 1| 3D rendering of a hippocampal MFB.** Volume segmentation (blue) and MFB-surface area occupied by pSCRs (white) of the enlarged MFB in Fig. 2b. 3D-reconstruction from volumetric, near-isotropically resolved (*z*-STED) coCATS data acquired in the CA3 *stratum lucidum* of an adult, coCATS *in vivo* microinjected and perfusion-fixed mouse.

**Supplementary Video 2| CoCATS volume and MFB-segmentations in the CA3 *stratum lucidum*.** Fly-through (*xz*-view) of the volume in Fig. 2h, including BASSOON (magenta, confocal, N2V), SHANK2 (turquoise, *z*-STED, N2V) and coCATS (grey, *z*-STED, N2V), as well as 10 segmented MFBs. MFB surface areas occupied by pSCRs are indicated in white. Step size: 50 nm (537 optical sections corresponding to 26.8 μm). For visualization, contrast limited adaptive histogram equalization (CLAHE, ImageJ) was used to account for differential photobleaching.

**Supplementary Video 3| CoCATS volume in hippocampal neuropil.** Fly-through of coCATS volume in Fig. 3a (*xy*-view, *z*-STED, N2V), acquired in the neuropil of an organotypic hippocampal slice. Step size: 50 nm (220 optical sections corresponding to 11 μm).

**Supplementary Video 4| Reconstructing the local input field of a CA3 pyramidal neuron from volumetric coCATS data.** Fly-through of coCATS imaging volume (grey, *z*-STED, N2V) with intracellular CA3 PN label (yellow, confocal) (*xy*-view, 160 optical sections corresponding to 8 μm). 3D-renderings of the CA3 pyramidal neuron proximal dendrite (gold) and 57 synaptically connected structures (multi-colored) reconstructed from coCATS data are shown, as displayed in Fig. 3h. Connected structures were identified via presence of pSCRs (white).

**Supplementary Video 5| Reconstruction of the local output structure of a mossy fiber in the DG hilus.** Volume rendering of a piece of DG granule cell axon (mossy fiber, orange) with three MFBs and their corresponding post-synaptic structures (green), identified through the presence of pSCRs (white), as displayed in Fig. 4d.

**Supplementary Video 6| Reconstruction of the local output structure of a single MFB in the CA3 *stratum lucidum*.** Volume rendering of an MFB (orange), including its axon and filopodial extensions, and 9 post-synaptic structures (turquoise/blue) identified through the presence of pSCRs (white), as displayed in Fig. 4e.

**Supplementary Video 7| coCATS volume in a human cerebral organoid.** Fly-through along *z*-direction and volume rendering of a piece of cerebral organoid (29.9 x 22.9 x 8.7 $\mu m^3$, Fig. 6c), coCATS labeled and imaged at near-isotropic super-resolution by STED microscopy, using contrast limited adaptive histogram equalization (CLAHE) for visualization.
